# ZDHHC17–Mediated Palmitoylation of Hepatitis E Virus ORF3 Protein Regulates Vectorial Trafficking in Polarized Epithelial Cells

**DOI:** 10.64898/2026.06.16.732616

**Authors:** Patrick Bröscky, Charlotte C. Syren, Di Ge, Leonid Kostrykin, Antonio Piras, Zoé Engels, Andrew Freistaedter, Huanting Chi, Paula Jordan, Charlotta Funaya, Karl Rohr, Sandhya P. Tiwari, Thibault Tubiana, Andreas Pichlmair, Viet Loan Dao Thi

**Affiliations:** Heidelberg University, Medical Faculty Heidelberg, Center of Infectious Diseases, Virology, Center for Integrative Infectious Disease Research, Heidelberg, Germany; Heidelberg University, Medical Faculty Heidelberg, Center for Infectious Diseases, Molecular Virology, Center for Integrative Infectious Disease Research, Heidelberg, Germany; Technical University of Munich, School of Medicine, Institute of Virology, 81675 Munich, Germany; Biomedical Computer Vision Group, BioQuant, IPMB, Heidelberg University, 69120 Heidelberg, Germany; 5Heidelberg University, Electron Microscopy Core Facility, 69120 Heidelberg, Germany; Institute for Protein Research, The University of Osaka, 3-2 Yamadaoka, Suita, Osaka, Japan; Institute for Integrative Biology of the Cell (I2BC), Université Paris-Saclay, CEA, CNRS, Gif-sur-Yvette, France; German Centre for Infection Research (DZIF), Partner Site Heidelberg, Heidelberg, Germany

**Keywords:** hepatitis E virus, open reading frame 3 protein, polarized epithelial cells, palmitoyl transferase ZDHHC17

## Abstract

The hepatitis E virus (HEV) is a leading cause of acute hepatitis worldwide and is transmitted enterically along the gut–liver axis. Epithelial cell polarity in the gut and liver plays a critical role in HEV transmission. In intestinal epithelial cells, HEV enters through the apical membrane and is released basolaterally to access the bloodstream, whereas in hepatocytes, the virus enters basolaterally and is secreted apically into the bile duct. In this study, we sought to identify the viral and host determinants governing directional HEV secretion in both tissue types. Using polarized intestinal and hepatocyte models, we found that the small phosphoprotein ORF3, which is essential for progeny secretion, localizes predominantly to the apical membrane, in contrast to the ORF2 capsid protein. We further identified the palmitoyltransferase ZDHHC17 as a specific ORF3 interactor that mediates its apical localization and promotes HEV progeny release. Using automated cell segmentation and quantitative image analysis, we screened a panel of ORF3 mutants and found that positively charged residues within the N-terminus regulate membrane association. In addition, we identified a conserved PIFIQP motif that mediates the critical interaction with the ankyrin repeat domain (ARD) of ZDHHC17. Leveraging this interaction, we generated high-confidence structural models of ORF3 in complex with the ZDHHC17 ARD using AlphaFold. These models, further validated by molecular dynamics simulations, revealed additional ORF3 residues involved in the interaction. Collectively, our findings define key mechanisms underlying directional HEV release and provide broader insights into trafficking processes in polarized epithelial cells.

## Introduction

Hepatitis E virus (HEV) is a major cause of acute hepatitis worldwide and belongs to the *Hepeviridae* family^1^. Human-infecting HEV genotypes are classified within the *Paslahepevirus balayani* species, which comprises eight genotypes (HEV-1 to HEV-8)^2^. HEV-1 and HEV-2, are restricted to humans and transmitted via the fecal–oral route, mainly through contaminated water. Although infection is typically self-limiting in healthy individuals, HEV-1 and HEV-2 are associated with high mortality rates in pregnant women^3^. In contrast, HEV-3 and HEV-4 are zoonotic and are primarily transmitted to humans through consumption of contaminated meat products, e.g. from domestic pigs or wild boar^4^. These genotypes are most prevalent in industrialized countries. While the majority of HEV-3 and HEV-4 infections are asymptomatic, they can become chronic in immunosuppressed individuals, who have an elevated risk of developing liver cirrhosis, and consequently, liver failure^5^.

The positive-sense, single-stranded HEV RNA genome of approximately 7.2 kilobases encodes at least three open reading frames (ORFs). ORF1 is thought to be translated as a polyprotein and encodes the nonstructural proteins required for viral replication (reviewed in^6^). ORF2 encodes the capsid protein and plays a critical role in counteracting innate and adaptive host immune responses (reviewed in^7^). ORF3 is a small phosphoprotein of approximately 113 amino acids that is proposed to function as an ion channel^8^. Phosphorylation at serine 80 is thought to mediate interaction with the capsid protein ORF2^9^. At its N-terminus, eight cysteine residues undergo palmitoylation, which is essential for membrane association of ORF3^10^. One of the most prominent roles of ORF3 is the secretion of HEV progenies^11^. Through its conserved proline–serine–alanine–proline (PSAP) motif, ORF3 interacts with the endosomal sorting complexes required for transport (ESCRT) protein TSG101^12^, thereby facilitating virus budding into multivesicular bodies (MVBs) in a non-cytolytic manner. During this process, and despite the absence of viral glycoproteins, HEV particles acquire a host cell–derived membrane envelope that is decorated with ORF3^13^. Consequently, HEV particles released into the cell culture supernatant or circulating in the blood of infected patients are quasi-enveloped^14^. Although this envelope, derived from the trans-Golgi network (TGN) membrane^15^, reduces cell attachment and entry, it protects viral particles from neutralizing antibodies, likely facilitating spread within the infected host. In contrast, HEV shed in feces or present in HEV-replicating cells are non-enveloped, highly stable, and infectious, enabling the transmission to new hosts^16^.

Epithelial cells are polarized in order to maintain their barrier function while enabling transport functions. To this end, they have distinct membranes: On the one hand, the apical membrane, facing the outside environment and on the other hand, the basolateral membrane, facing neighboring cells and underlying tissue. HEV uniquely depends on tissue epithelial cell polarity for its enteric transmission. Following entry via the gastrointestinal tract, HEV is thought to infect the intestinal epithelium through the apical membrane and enters the bloodstream through release from the basolateral membrane into the bloodstream. Once in circulation, the virus reaches the liver, where it enters hepatocytes via the basolateral membrane and is primarily released from the apical membrane into the bile duct^16^. When transported with the bile back to the intestine, the quasi-envelope is removed and HEV is ultimately shed in feces, enabling transmission to new hosts. Until now, knowledge of how the virus hijacks the polarized trafficking machinery of intestinal epithelial cells and hepatocytes has been lacking.

Recent studies have shown that the ORF3 protein is required for apical HEV progeny release from polarized hepatocytes and liver organoids^17–19^. In agreement, ORF3 was mainly located close to the apical membrane and within the bile canaliculi of human hepatocytes in a human-liver chimeric mouse model^20^. Surprisingly, we and others have shown a predominantly apical release of HEV progeny from polarized intestinal epithelial cells or intestinal organoids^21,22^, but the viral determinant governing this process was not identified.

In the present study, we confirm ORF3 to be the principal viral determinant governing apical HEV trafficking in polarized hepatocytes and extend this finding to polarized intestinal epithelial cells, thereby strongly supporting the intestine as a major site of HEV shedding during infection. We further identified the palmitoyltransferase ZDHHC17 as a key mediator of ORF3 palmitoylation and its role in apical HEV progeny release. Through systematic screening of mutants targeting the charged residues within ORF3, we uncovered novel sequence motifs that regulate its palmitoylation and/or mediates its interaction with ZDHHC17. We employed AlphaFold modeling and molecular dynamics simulations to gain additional mechanistic insights into the ORF3–ZDHHC17 interaction. Collectively, these findings reveal previously unrecognized viral and host determinants governing HEV release and provide new insights into vectorial trafficking in polarized epithelial cells.

## Results

### ORF3, but not the ORF2 protein, is mainly apically localized in polarized intestinal and liver epithelial cells

In order to identify the structural determinant governing vectorial release of HEV in either tissue context, we first investigated the subcellular localization of ORF2 and ORF3 proteins in hepatocytes and intestinal epithelial cells infected with the HEV-3 Kernow-C1/p6 strain. Notably, whereas intestinal epithelial cells typically exhibit a 2-dimensional (2D) columnar polarization, with an apical membrane facing the lumen and the basolateral side facing the bloodstream, hepatocytes display a more complex, 3-dimensional (3D) architecture with multiple apical and basolateral membrane domains *in vivo* (reviewed in^23^, Fig. 1A).

**Figure 1.**
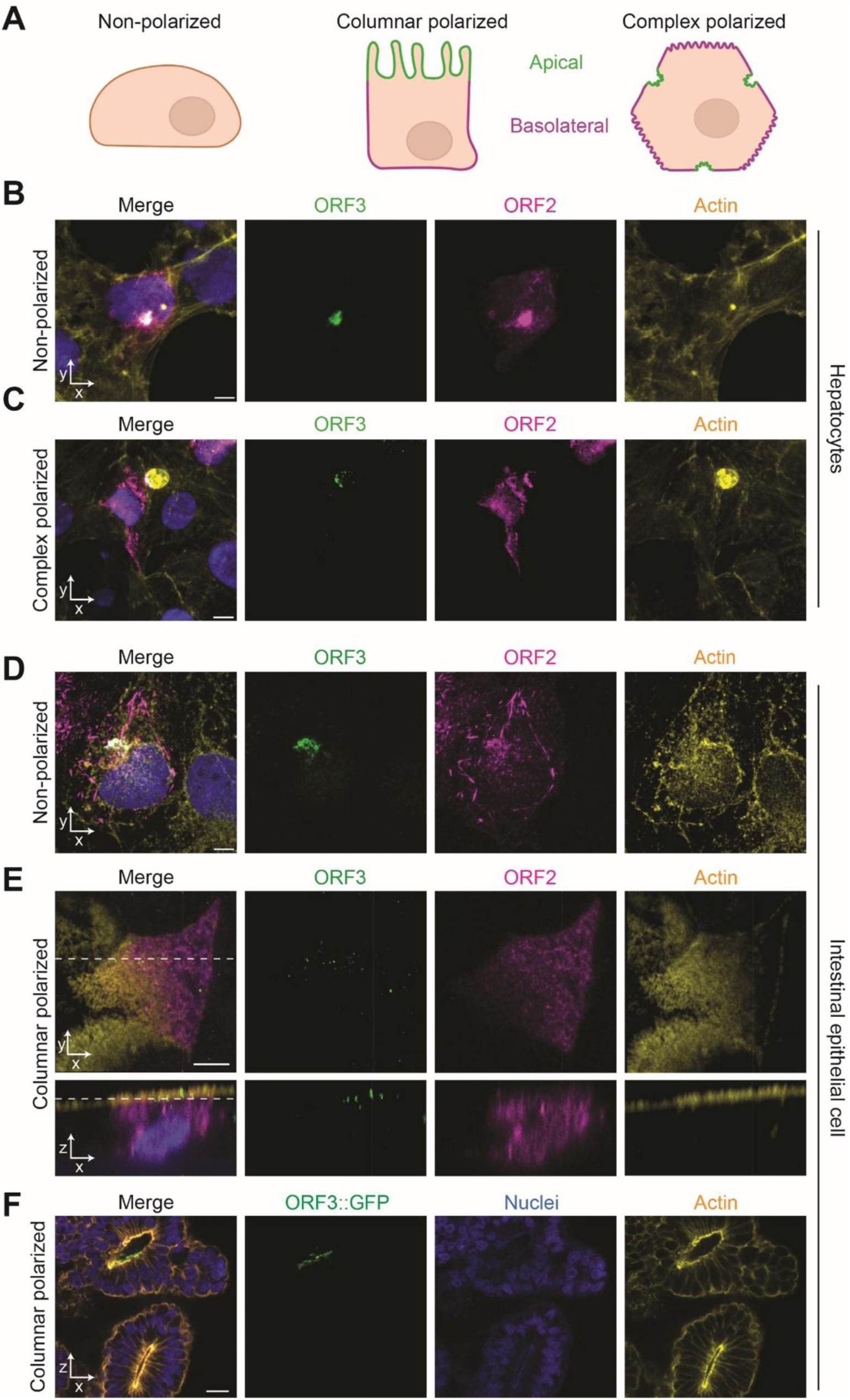
ORF3, but not ORF2 protein, is mainly localized apically in polarized intestinal and liver epithelial cells. **(A)** Schematic representation of non-polarized, columnar polarized and complex polarized cells. Apical (green) and basolateral (magenta) areas are indicated. Created in BioRender. Freistaedter, A. (2026) https://BioRender.com/i6730i9 **(B-E)** Hepatoma cells, including (B) non-polarized S10-3 and (C) complex polarized HepaRG, and intestinal epithelial cells, including (D) non-polarized, and (E) polarized Caco-2, were infected with HEV (MOI = 1 FFU/cell). 7 days post-infection, cells were fixed and stained for ORF2 (magenta), ORF3 (green), nuclei (blue) and actin (yellow). Images ((B-D) maximum projections, (E-F) single slice) were taken with a Zeiss Airyscan 2 LSM 900. (B-D) XY view, (E) XZ view. Scale bar = 5 µm. Images are representative of three independent experiments. **(F)** Intestinal organoids were transduced with ORF3::GFP, fixed after 48 h, and imaged. Scale bar = 200 µm.

First, we employed hepatocellular model systems that recapitulate cell polarity at increasing levels of complexity. In non-polarized hepatoma S10-3 cells infected with HEV, ORF2 was distributed throughout the cytoplasm and formed fibrillar structures, whereas ORF3 localized predominantly to a confined perinuclear region, consistent with previous reports^24^. Notably, a fraction of ORF2 colocalized with ORF3 within this perinuclear compartment. (Fig. 1B). Then, we infected columnar polarized human induced pluripotent stem cell (hiPSC)-derived hepatocyte-like cells (HLCs)^25^. Similar to non-polarized hepatocytes, ORF2 was detected throughout the cell body, while ORF3 localized almost exclusively to the apical membrane in polarized HLCs (Suppl. Fig. 1). Finally, we used bipotent HepaRG cells, which upon differentiation acquire a complex 3D polarization characterized by multiple apical and basolateral membrane domains, and the formation of functional biliary canaliculi-like structures^26^. In HEV-infected differentiated HepaRG cells, ORF2 again showed a broad intracellular distribution, whereas ORF3 localized predominantly to apical membranes, as evidenced by strong co-localization with F-actin or MRP2, which is enriched at the apical region of polarized hepatocytes (Fig. 1C, Suppl. Fig. 2)^27^.

Next, we investigated the localization of ORF2 and ORF3 in HEV-infected intestinal epithelial cells. In non-polarized intestinal Caco-2 cells, ORF2 and ORF3 showed a similar distribution to that observed in non-polarized hepatoma cells, with ORF2 present throughout the cells while ORF3 was confined to a defined perinuclear region (Fig. 1D). We then infected Caco-2 cells cultured on Transwell inserts, where they spontaneously develop a columnar polarized epithelial architecture^28^. In these 2D-polarized cells, consistent with observations in 2D-polarized HLCs and differentiated HepaRG cells, only ORF3 accumulated specifically at the apical membrane (Fig. 1E). Finally, we confirmed the exclusive localization of ectopic ORF3::GFP on the apical membrane of hiPSC-derived intestinal organoids facing their lumen, further supporting a conserved apical targeting in polarized intestinal epithelium (Fig. 1F)^22,29^.

Altogether, these results show that the HEV structural protein ORF3, but not ORF2, was mainly localized apically in both polarized intestinal cells and hepatocytes. With its established role in HEV secretion and previous reports describing preferential apical HEV release from polarized enterocytes^17,22^, our findings support ORF3 as a key determinant of directional HEV progeny release.

### ZDHHC17 mediates ORF3 palmitoylation to control its apical localization and HEV progeny release

To identify potential ORF3-interacting host factors involved in its apical trafficking and directional progeny release, we performed an ORF3 interactome screen. For this purpose, we ectopically expressed ORF3 in hepatoma HepG2/C3A cells as an N-terminal fusion protein with an HA-tag, alongside a mEGFP control. We immunoprecipitated ORF3::HA from cell lysates and identified associated proteins by mass spectrometry.

As shown in Fig. 2A, this approach identified numerous components of the vesicle sorting machinery, such as VPS28 or VPS37C. Among the strongest candidates was the ESCRT protein TSG101, which has previously been reported to interact with ORF3^30^. Importantly, we also identified ZDHHC17, a member of the aspartate–histidine–histidine–cysteine (DHHC) family of S-acyltransferases. This family spans at least 23 integral membrane enzymes called palmitoyl acyl transferases (designated ZDHHC 1–23) that mediate S-palmitoylation on cysteines. All members of the ZDHHC family share a conserved DHHC catalytic motif, located within a cysteine-rich domain that faces the cytosol and that is essential for enzymatic activity. Other important domains include transmembrane, SH3, PDZ binding, and ankyrin repeat domains (ARD)^31^.

**Figure 2.**
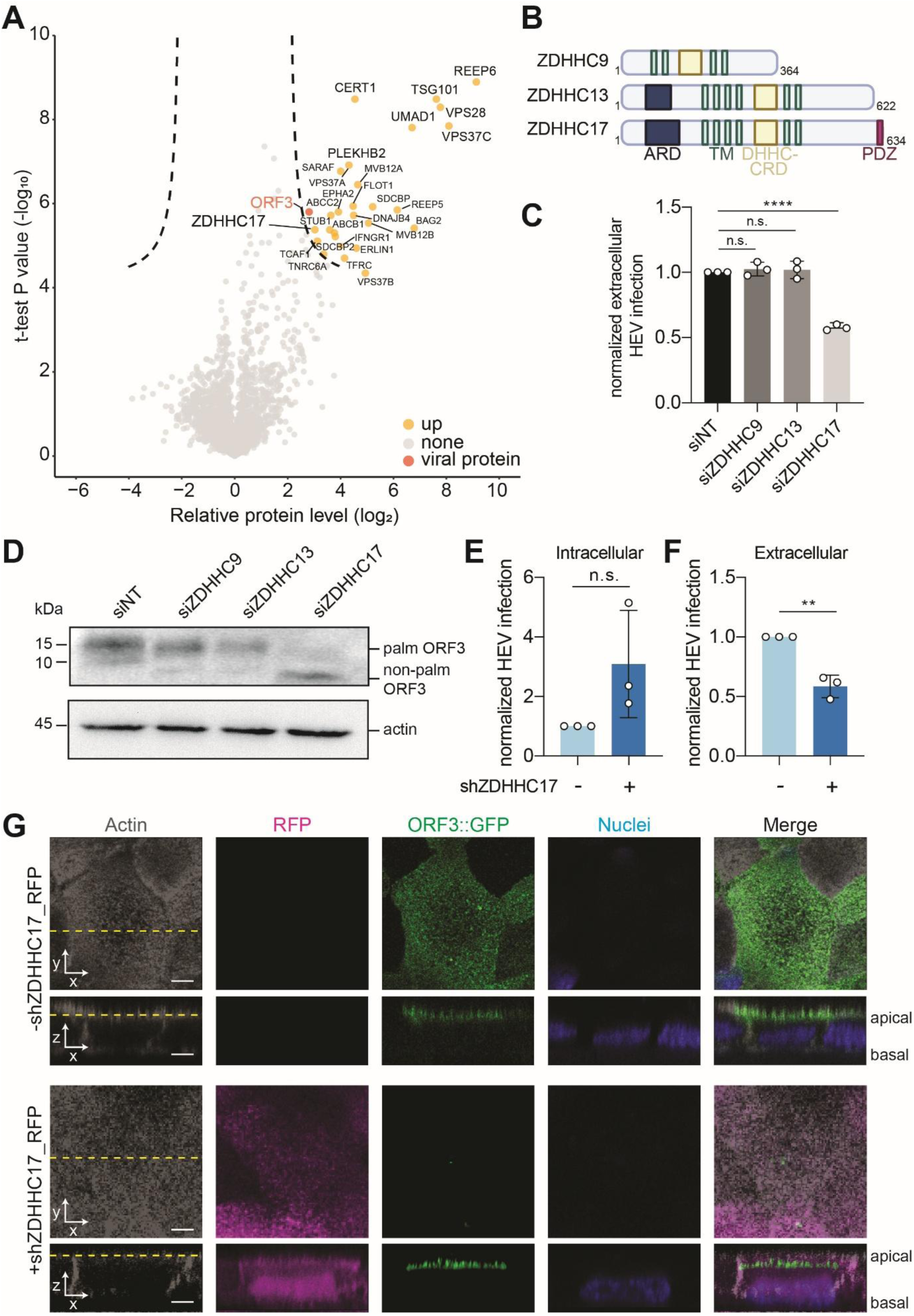
ZDHHC17 mediates ORF3 palmitoylation to control its apical localization and HEV progeny release. **(A)** HepG2/C3A cells expressing ORF3::HA or mEGFP were subjected to Co-IP followed by mass spectrometry. Volcano plot showing proteins identified in the mass spectrometry screen. Each point represents a detected protein, plotted according to log2 relative protein abundance and statistical significance (t-test, p-value, -log10). The thresholds for significance and relative protein level are indicated by dashed lines. Viral proteins (orange) and upregulated host proteins (yellow) are highlighted. **(B)** Scheme of protein domains in ZDHHC9, 13, and 17; ARD = Ankryin repeat domain, TM = transmembrane domain, DHHC-CRD = catalytic site domain, PDZ = protein-protein interaction module. Created in BioRender. Dao Thi, V. L. (2026) https://BioRender.com/ngauquj **(C)** Caco-2 cells electroporated with HEV Kernow-C1/p6 RNA were transfected 5 days post-electroporation with respective siRNAs (100 nM). Supernatant between day (d) 2 and 4 posttransfection was collected and titrated on HepG2/C3A. Infected HepG2/C3A cells were fixed 5 d postinfection and stained for ORF2 capsid protein to assess FFUs. Samples were normalized to siNT. n = 3 biological replicates. **(D)** Caco-2 cells expressing ORF3 were transfected with siNT, siZDHHC9, siZDHHC13 or siZDHHC17 (100 nM) and lysates were harvested 2 d later for WB analysis. **(E, F)** Caco-2 cells, expressing an dox-inducible plasmid with an shRNA against ZDHHC17, were electroporated with HEV Kernow-c1/p6 RNA. 5 d later, the cells were treated with dox to induce shZDHHC17 expression. 7 d post-electroporation, the medium was replaced and 2 d later, virus from cell lysates (intracellular) and supernatant (extracellular) were harvested and titrated on HepG2/C3A cells. FFUs were assessed as described in (C). (C, E, F) n = 3 biological replicates. Significance by unpaired t test:** p < 0.01, **** p < 0.0001, n.s. not significant. **(G)** Confocal images of Caco-2 cells stably expressing a bicistronic dox-inducible RFP/shZDHHC17 (from E, F). During polarization the cells were dox-induced. When they reached a TEER of 1000 Ω×cm^2^, cells were transduced with lentivirus carrying an expression construct for ORF3::GFP. After 24 h, cells were fixed, stained for actin, and imaged using a Zeiss Airyscan 2 LSM 900 microscope. Scale bar = 5 µm. Images are representative of three independent experiments.

To validate the role of ZDHHC17 in HEV progeny release from intestinal cells, we performed siRNA-mediated knockdown of ZDHHC17 alongside ZDHHC13, the only other ZDHHC family member containing an ARD, which is implicated in protein-protein interactions (Fig. 2B). We also knocked down ZDHHC9, a distantly related family member that lacks the ARD. Intestinal Caco-2 cells were electroporated with *in vitro*-transcribed Kernow-C1/p6 RNA, followed by transfection with siRNAs to downregulate the respective transferase. We assessed the release of infectious particles into the supernatant by titration on naïve HepG2/C3A cells and quantified ORF2-positive foci forming units.

Compared to the non-targeting (NT) control, knockdown of ZDHHC9 and ZDHHC13 had no detectable effect, whereas downregulation of ZDHHC17 reduced the release of infectious HEV progeny by approximately 50% (Fig. 2C). This reduction correlated with the efficiency of ZDHHC17 mRNA knockdown, as assessed by qRT-PCR (Suppl. Fig. 3A). Western blot analysis of transfected cell lysates further revealed that only ZDHHC17 knockdown, but not that of ZDHHC9 or ZDHHC13, abolished ORF3 palmitoylation (Fig. 2D). Together, these findings indicate that ZDHHC17 specifically, rather than other palmitoyl acyltransferases, is required for the trafficking and secretion of infectious HEV particles.

To achieve a more efficient downregulation of ZDHHC17, we generated a stable Caco-2 cell line using a selectable lentiviral system expressing a doxycycline (dox)-inducible shRNA targeting ZDHHC17 (Caco-2_shZDHHC17), which is bicistronically linked to a red fluorescent protein (RFP). When compared to siRNA-mediated knockdown, this approach resulted in a more pronounced reduction of ZDHHC17 mRNA levels (Suppl. Fig. 3B).

We electroporated Caco-2_shZDHHC17 cells with *in vitro*-transcribed HEV Kernow-C1/p6 RNA, treated the cells with dox, and harvested intra- and extracellular viral progenies for titration. While the downregulation of ZDHHC17 led to a non-significant accumulation of intracellular infectious particles (Fig. 2E), it significantly reduced the release of extracellular progeny (Fig. 2F).

Since ORF3 localizes to the apical region in both polarized intestinal epithelial cells and hepatocytes, and the columnar polarization of intestinal epithelial cells allows a clear and rapid distinction between apical and basolateral membranes, we used (columnar polarized) Caco-2 cells for the remainder of the study. Caco-2_shZDHHC17 cells were cultured on Transwell inserts until polarization was reached, as indicated by a transepithelial electrical resistance (TEER) of >1000 Ω×cm² ^32^. Once polarized, shRNA expression was induced by dox-treatment. Two days later, cells were transduced to express a GFP tagged ORF3 (ORF3::GFP) to assess the role of ZDHHC17 in apical ORF3 localization. Importantly, the GFP-tag did not affect ORF3 apical localization (Suppl. Fig. 4A) and still allowed its secretion via multivesicular bodies (MVBs), although at lower levels than untagged ORF3 (Suppl. Fig. 4B, 4C).

24 hours later, cells were fixed and z-stack images were acquired by confocal microscopy. As shown in Fig. 2G, in polarized Caco-2 cells expressing shZDHHC17 (as indicated by RFP expression), ORF3::GFP failed to fully reach the apical membrane and instead accumulated beneath it. In contrast, in non-dox induced RFP-negative cells, ORF3::GFP was predominantly localized at the apical side. Taken together, these results identified ZDHHC17 as the specific palmitoyltransferase responsible for ORF3 palmitoylation, thereby regulating its apical trafficking, and the release of infectious HEV progeny.

### Mutational screening of charged residues in ORF3 reveals novel motifs critical for its apical localization in polarized cells

Viral proteins hijack the cellular trafficking machinery to navigate in polarized cells and reach specific subcellular regions (reviewed in^33^). To access this machinery, viruses often encode residues that mediate binding to host trafficking factors and/or acquire post-translational modifications that serve as sorting signals^34,35^. Thus, we next performed an alanine-scanning mutagenesis screen across ORF3 to assess which residues could be involved in its apical localization in polarized cells.

To analyze the impact of mutations on apical ORF3 localization, we developed an automated image analysis pipeline that enables quantitative assessment of fluorescence signals in 2D-columnar polarized cells (Fig. 3A). To this end, we stained polarized Caco-2 cells with MemBrite®-Fix, which is a specific dye for plasma membranes, and acquired **∼**130 z-stack images (spacing of 0.13 µm) per cell on a confocal microscope.

**Figure 3.**
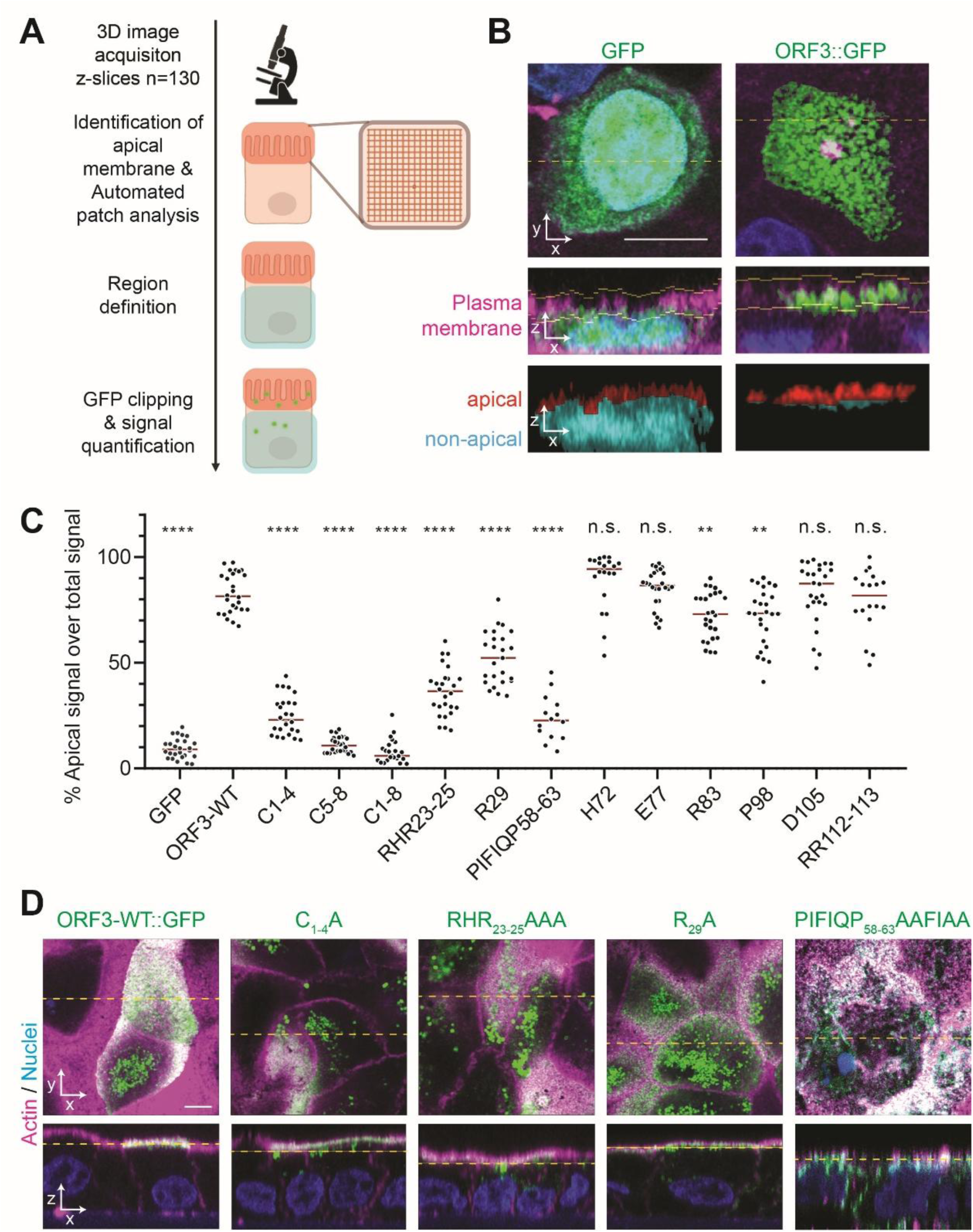
Mutational screening of charged residues in ORF3 reveals novel motifs critical for its apical localization in polarized cells. **(A)** Overview of the automated image analysis pipeline for quantifying GFP apical signal in polarized cells. Created in BioRender. Dao Thi, V. L. (2026) https://BioRender.com/pkbq0hh **(B)** Example images to illustrate the workflow shown in (A) of polarized Caco-2 cells transduced to express GFP or ORF3::GFP (green). 5 d posttransduction, cells were fixed, stained for nuclei (blue) and the plasma membrane (magenta) and imaged with a Zeiss Airyscan 2 LSM 900. The yellow line indicates the position of the XZ slice shown in the second row. Plasma membrane staining with MemBrite (magenta) was used to define the apical region, highlighted by the yellow boundaries. The third row shows the final segmentation of the GFP signal into apical (red) and non-apical (cyan) regions. Scale bar = 10 µm. **(C)** Polarized Caco-2 cells were transduced to express GFP or ORF3::GFP variants and imaged as described in (B). The percentage of apical GFP over the total apical and non-apical signal was quantified by the automated image analysis pipeline shown in (A). n = biological replicates. Red line: Median. Significance by ordinary one-way ANOVA: ** p < 0.01, **** p < 0.0001, n.s. not significant. **(D)** Representative confocal images of Caco-2 cells transduced to express ORF3::GFP, C_1-4_A, RHR_23-25_AAA, R_29_A, PIFIQP_58-63_AAFIAA (green). Cells were fixed and stained for actin (magenta) and nuclei (blue). Top row: XY, bottom row XZ view. Scale bar = 5 µm.

Region of interest (ROI) selection was performed by maximum-intensity projection of the GFP channel of the 3D image along the z-axis after exclusion of the lowest five z-slices, thresholding the resulting maximum-intensity projection image using Otsu’s method^36^, morphological hole filling, and identifying the connected components. If the largest connected component covered 5–20% of the maximum-intensity projection image, the bounding box of this component was expanded symmetrically by 25% along the x-axis and the y-axis, and then selected as ROI. The apical region was identified based on its enrichment of upper membrane signals. To localize the apical region, the membrane channel was partitioned into 12 × 12 pixel patches in the xy-plane. For each patch, membrane intensity profiles along the z-axis were computed and the uppermost z-position satisfying one of two conditions was selected as the top border of the apical region: (1) The membrane intensity at this z-position and the next two z-positions below exceeded a threshold, or (2) the membrane intensity profile at this z-position and the next z-position below exceeded a threshold, and also the intensity at the next z-position below was higher than that at the next z-position above. The threshold was determined adaptively by using the mean value of the membrane intensity profile, or half of the mean if the intensity distribution was right-skewed. The apical region was then defined as a band below the top border, with the non-apical region located below it (Fig. 3B). To reduce image noise, GFP signals were clipped at the 99.9^th^ percentile of the GFP intensity distribution, smoothed using a 3D Gaussian filter, and the background signal was excluded by Otsu thresholding^36^. The GFP intensities were subsequently quantified within apical and non-apical regions, and the percentage of apical GFP over the total apical and non-apical signal was computed.

To screen for residues involved in apical ORF3 localization, columnar polarized Caco-2 cells were ectopically transduced with lentiviral constructs encoding GFP, GFP-tagged ORF3 wild type (WT), or a panel of GFP-tagged ORF3 mutants. In the GFP-tagged ORF3 mutants, charged amino acid residues were substituted with alanine (Fig. 3C). We additionally mutated the eight N-terminal cysteine residues previously reported to be palmitoylated, as well as the C-terminal PSAP motif required for interaction with TSG101. We further mutated the PIFIQP motif (residues 58–63) to AAFIAA; a motif that is highly conserved among *Paslahepevirus balayani* species members (Suppl. Fig. 5). This sequence corresponds to the previously described ΨβXXQP motif, which has been implicated in interactions of substrate proteins with ZDHHC17^37^. We transduced Caco-2 cells with respective ORF3::GFP encoding lentiviruses and seeded them on Transwells. After polarization, we stained the cells with MemBrite® Fix, fixed the cells, acquired z-stack images on a confocal microscope, and analyzed them using our quantitative image segmentation and analysis pipeline.

As shown in Fig. 2C, GFP alone exhibited only 10% apical localization, whereas 80% of WT ORF3::GFP was localized apically, almost exclusively co-localizing with the apical actin staining (exemplary images used for the pipeline in Suppl. Fig 6 and super-resolution images in Fig. 3D). Mutating either the first or the last four N-terminal cysteine residues reduced ORF3::GFP apical localization to 30% and 10%, respectively. While some of the ORF3 C1-4A mutant still co-localized with apical actin, a major fraction was clearly retained inside the cell (Fig. 3D). When all cysteine residues were substituted with alanine, we detected only 10% of the ORF3::GFP signal apically, comparable to GFP alone.

Interestingly, mutation of two clusters of charged residues located near the cysteine residues (RHR23–25AAA and R29A) reduced apical localization of ORF3::GFP by approximately 40%. These mutants appeared to be retained beneath the apical actin staining (Fig. 3D). All other mutations in the ORF3 C-terminus, including the PSAP motif, did not significantly affect apical localization. Interestingly, the AAFIAA mutant exhibited 25% apical localization. Furthermore, as shown in Fig. 3D, this mutant partially redistributed to the basolateral membrane of polarized Caco-2 cells, with a fraction also detected below apical actin.

### The ΨβXXQP motif in ORF3 mediates its interaction with ZDHHC17 and HEV progeny egress

Having identified N-terminal charged residues and a putative C-terminal ZDHHC17-interacting motif, we next assessed the impact of these mutations on the interaction with ZDHHC17. To this end, HA-tagged ZDHHC17 was co-expressed with GFP-tagged ORF3-WT or mutants in HEK293T cells. Following GFP-Trap pulldown and Western blot analysis, ORF3::GFP WT, as well as the RHR23–25AAA and R29A mutants, retained interaction with ZDHHC17 (Fig. 4A). This interaction was further supported by immunofluorescence analysis, which revealed co-localization of ORF3 with ZDHHC17 in HEV-replicating cells (Suppl. Fig. 7A). In contrast, the ORF3::GFP AAFIAA mutant failed to interact with ZDHHC17 (Fig. 4A).

**Figure 4.**
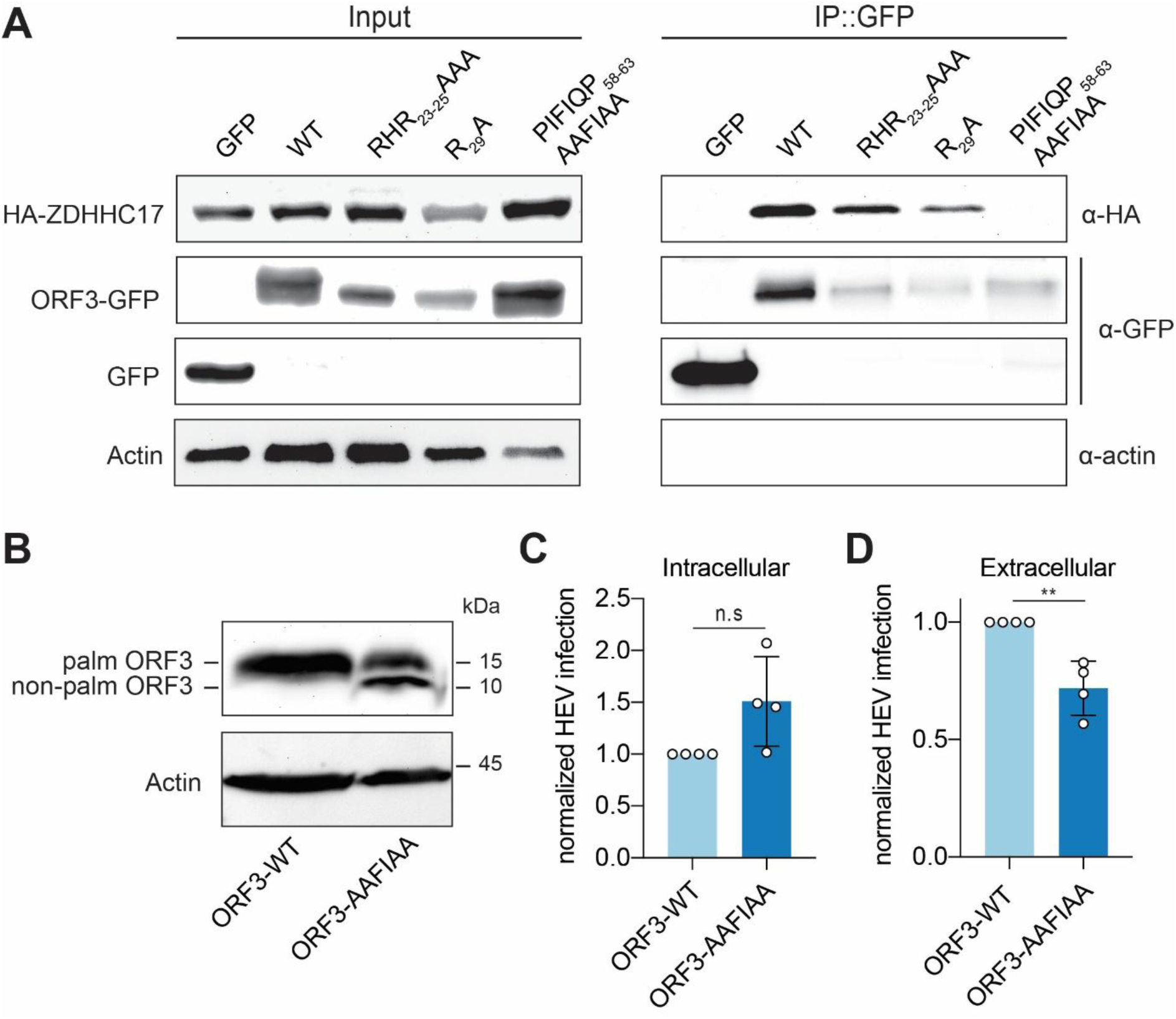
The ΨβXXQP motif in ORF3 mediates its interaction with ZDHHC17 and HEV progeny egress. **(A)** HEK293T cells were co-transfected with plasmids encoding ZDHHC17::HA and ORF3::GFP WT and derivatives and lysed 24 h post-transfection, followed by anti-GFP IP and WB analysis for ORF3 (α-GFP), ZDHHC17 (α-HA) and β-actin expression, using indicated antibodies. **(B)** WB analysis of Caco-2 cells stably expressing ORF3-WT or ORF3-AAFIAA using antibodies against ORF3 and β-actin. **(C+D)** ORF3 trans-complementation: Caco-2 cells stably expressing ORF3-WT or ORF3-AAFIAA were electroporated with ΔORF3 HEV Kernow-C1/p6 RNA. 5 days later media was changed. 2 days later HEV particles were harvested intra- and extracellular and titrated on HepG2/C3A cells. FFUs were assessed 5 days post-infection by staining against HEV ORF2 protein. n = 4 biological replicates. Significance by unpaired t test:** p < 0.01, n.s. not significant.

Importantly, all mutants exhibited a lower molecular weight compared to ORF3-WT (Fig. 4A). To more clearly distinguish between palmitoylated and non-palmitoylated ORF3 species, we also analyzed untagged ORF3 proteins. As shown in Fig. 4B, the ORF3-AAFIAA mutant was only partially palmitoylated compared to ORF3-WT, which we predominantly detected in its fully palmitoylated form. In contrast, the RHR23–25AAA and R29A mutants, despite retaining interaction with ZDHHC17, were primarily present in their non-palmitoylated form (Suppl. Fig. 7B). Importantly, substitution of the arginine residues with similarly positively charged lysins rescued ORF3 palmitoylation and apical localization (Suppl. Fig. 7C, 7D).

ORF2 and ORF3 are expressed from a bicistronic RNA and therefore cannot be analyzed independently in the viral genome. To further assess the functional relevance of the ORF3 PIFIQP motif in HEV progeny release, we established an ORF3 trans-complementation system. We electroporated in vitro-transcribed Kernow-C1/p6 genomes with a mutated ORF3 start codon into Caco-2 cells ectopically expressing either ORF3-WT or ORF3-AAFIAA. On day 7 post-electroporation, we harvested intracellular virions from cell lysates and extracellular virions from cell culture supernatants and titered them on HepG2/C3A cells. As shown in Fig. 4C and 4D, ectopic expression of ORF3-AAFIAA resulted in non-significant intracellular accumulation but significantly reduced extracellular release of infectious HEV progeny compared to ORF3-WT expression, confirming the importance of ZDHHC17-ORF3 interaction in the HEV life cycle.

### Structural modeling and molecular dynamics simulations define the molecular determinants of ORF3–ZDHHC17 interaction

Previous studies have described ORF3 as an intrinsically disordered protein^38^, which heavily impairs reliable modeling of its structure and protein-protein interactions. Despite this, we aimed to further characterize the molecular basis of the interaction between ORF3 and ZDHHC17 using AlphaFold. To this end, we generated full-length complex models using AlphaFold3, considering both palmitoylated and non-palmitoylated ORF3 together with the ZHHC17 ARD. Consistent with its intrinsically disordered nature, ORF3 exhibited globally low confidence scores (pLDDT, Suppl. Fig. 8A, 8B). Nevertheless, in models incorporating palmitoylation, the PIFIQP motif was more frequently positioned in proximity to the ZDHHC17 ARD, as reflected by a shorter average distance between the center of geometry (COG) of the ARD and the PIFIQP motif (19.8 Å ± 5.7 Å, Suppl. Fig. 8C) compared to non-palmitoylated models (31.1 Å ± 8.1 Å, Suppl. Fig. 8D). While these differences are modest, they supported the rationale for exploring alternative modeling strategies using truncated ORF3 constructs focused on the interaction region. This does not necessarily reflect a true stabilizing effect of lipidation, but rather the fact that AlphaFold performs better when more of the biological context is included.

In the model with the highest confidence score, palmitoylated cysteines consistently localized in proximity to the catalytic DHHC motif (residues 464–467), while the membrane insertion predicted using the PPM webserver, supported a coherent orientation in which the transmembrane region of ZDHHC17 is correctly embedded and palmitoylated ORF3 cysteines are positioned within the membrane plane (Fig. 5A)^39,40^. Interestingly, basic residues within the N-terminal region of ORF3 identified in our mutagenesis screen (RHR23–25 and R29) were oriented toward the membrane interface, suggesting a role in enhancing electrostatic interactions with the membrane (Fig. 5Ai). Together, these observations support a model in which palmitoylation anchors ORF3 to the membrane and spatially constrains it near the catalytic site of ZDHHC17 (Fig. 5Aii).

**Figure 5.**
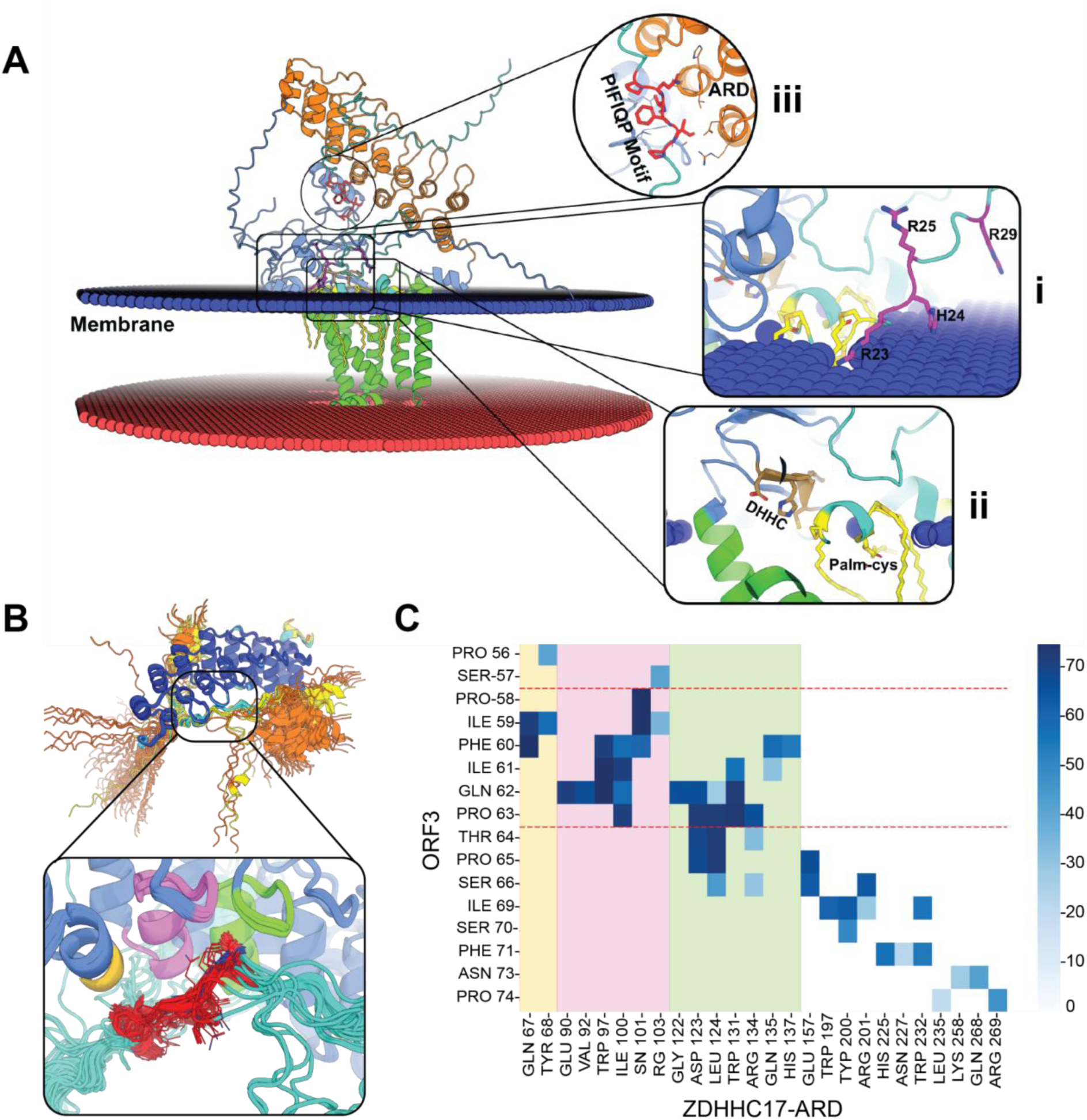
AlphaFold modeling of the ORF3_PIFIQP-ZDHHC17_ARD interaction reveals positioning of positive charged residues and consecutive positioning of the recognition sequence of ORF3 at the ARD of ZDHHC17. **(A)** AlphaFold3 model of the best-scoring ORF3–ZDHHC17 complex positioned relative to a lipid bilayer (blue, red). ZDHHC17 is shown embedded in the membrane, with the ankyrin repeat domain (ARD, orange) highlighted. ORF3 (orange) is represented as a largely disordered chain. Insets show (i) the orientation of basic residues (RHR23-25, R29; magenta) toward the membrane interface, (ii) the positioning of palmitoylated cysteines (yellow) near the DHHC catalytic motif (brown) and (iii) the proximity of the PIFIQP motif to the ARD. Membrane insertion was predicted using the PPM webserver. **(B)** Superposition of multiple models (from Sup. Fig. 7) demonstrating convergence of the PIFIQP motif onto a common binding region on the ARD surface. **(C)** Contact frequency map identifying key interacting residues between ORF3 and the ARD, revealing three main interaction regions.

Next, we investigated the interaction between the ORF3 PIFIQP motif and the ZDHHC17 ARD, as the ARD is known to mediate recognition of specific interaction motifs in substrate proteins. Using full-length ORF3, the PIFIQP motif was frequently positioned in proximity to the ZDHHC17 ARD (Fig. 5Aiii); however, the interaction exhibited low local confidence scores, precluding precise characterization of the interface. To improve prediction confidence, we focused on a truncated ORF3 segment spanning ±20 amino acids around the PIFIQP motif (residues 37–83). For ZDHHC17, only the ARD (residues 55–288) was included. Based on these confined regions, extensive AlphaFold2 sampling yielded highly confident predictions (pLDDT > 90) centered on the PIFIQP motif (Suppl. Fig. 9). As shown in Fig. 5B, in these models, the motif consistently occupied the same surface region of the ARD, indicating a reproducible binding mode rather than a model-specific arrangement.

We next performed an analysis (based on <4 Å distance cutoff) to define contacts, identifying three principal ARD regions that contribute to the interaction (Fig. 5C): a short helical segment (residues 67–68) and two helix–loop regions (residues 90–103 and 122–137). Key residues involved included TRP97, ILE100, ASN101, ASP123, LEU124, TRP131, ARG134, and GLN135. The interface is characterized by a combination of hydrophobic and polar contacts, with a pronounced contribution from hydrophobic residues forming a patch on the ARD surface. Importantly, residues immediately adjacent to the PIFIQP motif (THR64, PRO65, SER66, and ILE69) also contributed significantly to binding, indicating that the interaction extends beyond the core hexapeptide and involves a broader sequence context.

To assess the stability of the predicted interface, we performed molecular dynamics (MD) simulations using the same ORF3 fragment spanning residues 37–83. Four representative models were selected from extensive AlphaFold2 sampling to ensure robust statistics and capture conformational diversity within the truncated ORF3 construct.

Free-energy landscapes projected along principal components revealed heterogeneous behavior across these models: some conformations remained confined to narrow energy basins (models 0 and 1), whereas others sampled broader conformational space (models 10 and 16; Fig. 6A). Notably, we superposed structures on the ARD and performed principle component analysis on a minimal ORF3 segment encompassing the interaction interface (±4 residues around the PIFIQP motif), reducing noise to focus on the flexibility of this segment. Although the ARD was not restrained, it remained relatively stable throughout the simulations.

**Figure 6.**
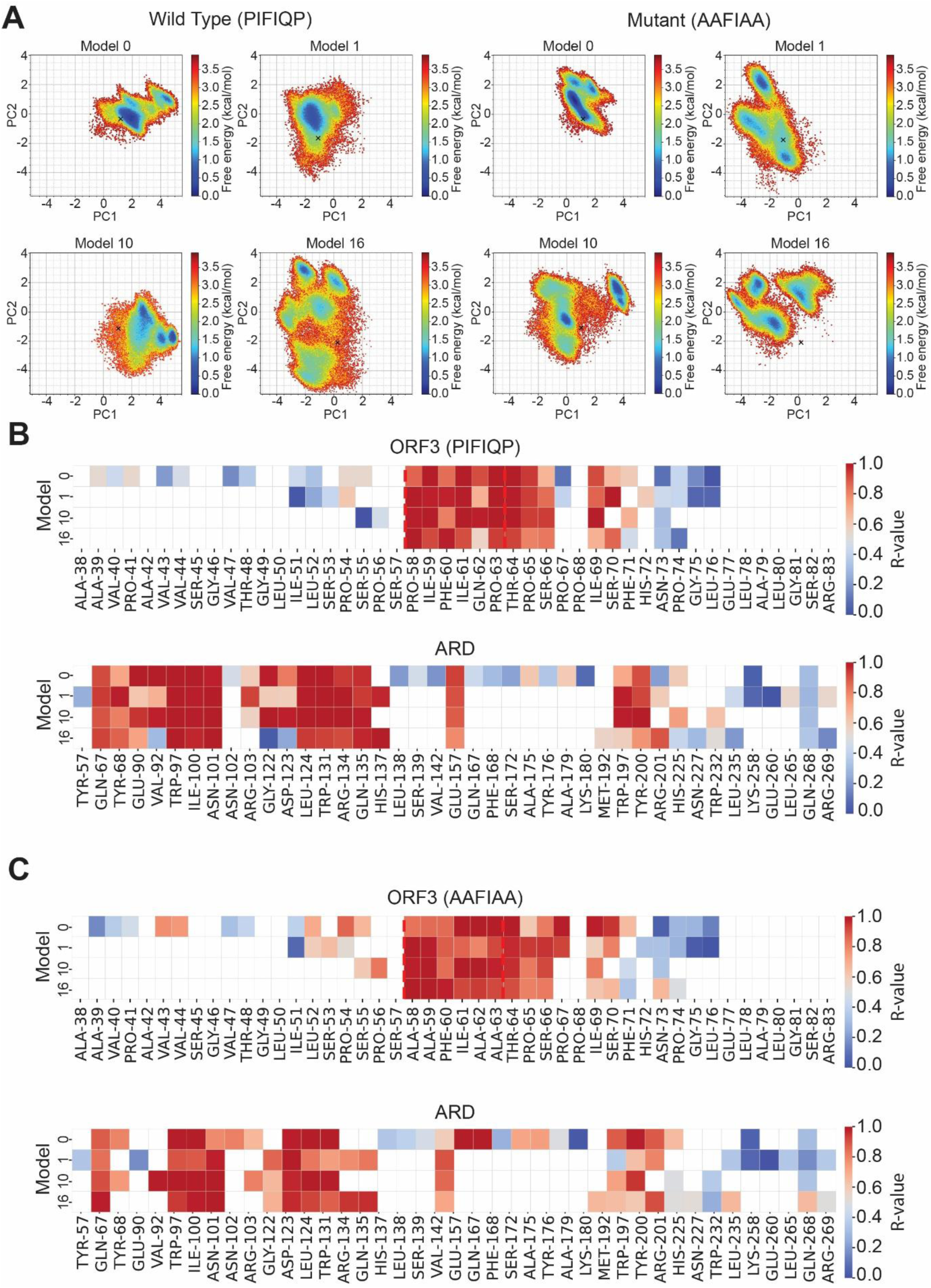
Conformational landscapes and contact stability analysis reveal an extended interaction interface between ORF3 and ZDHHC17. **(A)** Molecular dynamic simulations of ORF3 PIFIQP (± 4 residues up/downstream) together with the ARD of ZDHHC17. Free energy landscapes projected onto principal components (PC1, PC2) for representative models of the wild-type and AAFIAA variants. **(B, C)** Residue-wise contact persistence (R-values) for ORF3 and ARD residues across simulations, highlighting stable interactions centered on the wildtype, PIFIQP, motif (B) or the mutated, AAFIAA, motif (C) and its flanking residues.

Despite the observed variability among the four models, residue-wise contact stability (R-value^41^, calculated using a ≤5 Å distance cutoff) showed that the interaction between ORF3 and the ARD was consistently maintained (Fig. 6A). The same ARD residues retained high contact frequencies, supporting a stable binding interface once formed. In addition to the wild-type PIFIQP sequence, we simulated the AAFIAA mutant (both shown in Suppl. Vid. 1). This substitution disrupted GLN62 within the motif and reduced interaction density in the C-terminal region of the ARD (Fig. 6B, 6C), highlighting the importance of this residue for complex stabilization. The MD simulations further confirmed the involvement of key C-terminal ARD residues in interacting with the ORF3 PIFIQP motif (Fig. 6B, 6C), consistent with the AlphaFold2-based contact analysis (Fig. 5C).

Interestingly, residues flanking the canonical motif (THR64, PRO65, SER66) showed increased contact stability in the AAFIAA mutant, suggesting compensatory interactions. This behavior likely reflects the physicochemical properties of the interface: the ARD binding region forms a partially hydrophobic patch, and substitution with alanine residues does not strongly disrupt interactions once the complex is established. Consequently, mutations primarily modulate local interaction patterns rather than abolishing binding within the simulated timescale. This interpretation is consistent with the fact that all simulations were initiated from pre-formed wild-type complexes, thereby favoring retention of the bound state.

Overall, these results support a model in which basic residues within the N-terminal region facilitate membrane anchoring, thereby positioning ORF3 for interaction with ZDHHC17, while the ORF3 PIFIQP motif and its flanking residues mediate binding to the ZDHHC17 ARD.

## Discussion

The directional release of HEV particles along the gut–liver axis has previously suggested the existence of a cell type-specific trafficking machinery that differs between polarized hepatocytes and intestinal epithelial cells. In this study, we demonstrate that directional HEV trafficking toward apical membranes in polarized epithelial cells appears largely tissue-context independent. We identified ORF3 as the key structural viral determinant that requires lipidation by the palmitoyltransferase ZDHHC17. Previous studies showed that palmitoylation of ORF3 is important for membrane association^10^; here, we demonstrate that this modification also regulates its apical trafficking. An unbiased mutagenesis screen, combined with AlphaFold modeling and molecular dynamics simulations, further indicated that this process is governed by N-terminal basic residues that facilitate membrane anchoring, thereby likely positioning ORF3 in proximity to the catalytic site and enabling interaction with the ZDHHC17 ARD through its PIFIQP motif. Importantly, we show that this interaction is critical for efficient ORF3 palmitoylation, which in turn mediates apical trafficking of ORF3 and promotes HEV progeny release.

### Conserved apical HEV release from polarized epithelial cells along the gut-liver axis

Despite the distinct functions and polarization of epithelial cells in the liver and intestine, apical ORF3 trafficking appears to be conserved across these tissues. Consistent with this, enteric viruses are typically released from the apical membrane of the intestinal epithelium^42,43^, enabling transmission *via* the intestinal lumen and facilitating rapid, efficient spread between hosts, as observed, for example, for rotaviruses. In contrast, viruses targeting the liver are generally released from the basolateral surface of hepatocytes^44,45^, promoting systemic dissemination *via* the bloodstream.

Although we have used apical ORF3 trafficking as a proxy for apical release, our findings are consistent with recent reports demonstrating active infection of the gut and apical progeny release from intestinal cultures^17,22^. Together, these observations support the notion that the intestine represents a major site of HEV infection, particularly during acute, self-limiting infections, which are often asymptomatic or mild and lack classical hepatitis symptoms such as jaundice.

While the majority of HEV particles are released apically from polarized enterocytes, a fraction is also secreted basolaterally, likely as quasi-enveloped particles^17^. Although this pathway appears inefficient, it still permits viral spread to the liver and contributes to intra-host dissemination. Subsequent liver infection could, in turn, favor prolonged or even persistent infection, potentially due to the relatively immunotolerant hepatic environment^46^.

### ZDHHC17 is the specific enzyme mediating palmitoylation of ORF3 and thereby its apical trafficking

While polarized trafficking pathways in epithelial cells are generally well characterized, it remains poorly understood how viruses exploit these mechanisms to achieve directional release^47^. In particular, the molecular determinants that direct viral proteins into apical trafficking routes remain largely undefined.

Our findings indicate that palmitoylation is a key determinant in this process. This lipid modification promotes membrane association and influences partitioning into specific membrane microdomains, such as lipid rafts, which are linked to apical sorting via recycling endosomes^48–50^. Previous studies have shown that palmitoylation of ORF3 mediates membrane association and facilitates HEV progeny release^10^. Here, we extend these observations by demonstrating that palmitoylation also functions as a sorting signal that favors apical trafficking.

Future studies, for example, through biochemical isolation and characterization of lipid raft-associated ORF3 complexes in polarized cells, will be required to further dissect how lipid modifications contribute to the spatial organization of directional viral trafficking pathways. In addition, it will be important to determine the intracellular fate of non-palmitoylated ORF3, which is the predominant form either in the presence of downregulated ZDHHC17 (Fig. 2) or following mutation of the ORF3 PIFIQP motif (Fig. 4), and to identify the compartments to which it is redirected instead of trafficking to the apical membrane.

The specificity of ZDHHC17 in mediating ORF3 palmitoylation was unexpected, particularly given the close homology to ZDHHC13, which did not show a comparable effect in our system. However, substrate recognition within the ZDHHC family is not completely understood, and both overlapping and selective enzyme-substrate relationships have been reported^51^. In addition, specific interaction motifs within substrates can be recognized by dedicated domains such as the ARD of ZDHHC17, which mediates selective protein recruitment^52^. Importantly, MD analysis revealed that residues within AR domain 5—specifically TRP197, TYR200, and ARG201—also contribute to the ORF3–ZDHHC17 interaction. Although ZDHHC13 is generally homologous to ZDHHC17, these critical residues differ between the two proteins (Suppl. Fig. 10). This highlights that, despite overall redundancy within the ZDHHC family, individual enzymes can display pronounced substrate specificity, a feature that has been described for other viral proteins exploiting host palmitoylation machinery^53,54^. This specificity makes ZDHHC17 an attractive candidate for targeted antiviral intervention, as modulation of individual palmitoyltransferases is increasingly considered a viable therapeutic strategy^55^.

### Interaction with ZDHHC17 enables modeling of ORF3 structure

As ORF3 is predicted to be largely intrinsically disordered traditional structure-based modeling has been notoriously challenging and the structural basis of ORF3 function has remained elusive^38^. Here, by placing ORF3 in a more physiological context, including palmitoylation and membrane embedding, we obtained a high-confidence model using AlphaFold3 for the first time.

This model provided first structural insights into the ORF3–ZDHHC17 interaction despite the overall disorder of the protein. First, this model revealed that N-terminal basic residues adjacent to the critical cysteines do not appear to play a role in ZDHHC17 interaction *per se*, but are rather oriented toward the membrane interface, suggesting a role in enhancing electrostatic interactions with the membrane. This is consistent with previous reports showing that positively charged residues are often located near acylated cysteines in diverse cellular proteins^56^. Supporting this, substitution of arginines with lysins restored ORF3 palmitoylation and apical localization (Suppl. Fig. 7C, 7D) and RHR23–25AAA and R29A mutants retained their ability to interact with ZDHHC17 (Fig. 4A). We propose that these residues are rather essential for anchoring ORF3 into the membrane, thereby spatially constraining it near the catalytic site of ZDHHC17 to enable efficient palmitoylation.

In the full-length model, the ORF3 PIFIQP motif was positioned in proximity to the ZDHHC17 ARD; however, the interaction showed low local confidence scores, precluding precise definition of the interface. To address this, we focused on a truncated ORF3 sequence in complex with the ZDHHC17 ARD. Both AlphaFold modeling and MD simulations supported a specific interaction between these domains and identified additional critical residues. On ORF3, these findings extend the previously described ΨβXXQP motif to include THR64, PRO65, SER66, and ILE69, indicating that the interaction involves a broader sequence context beyond the core hexapeptide^37^. This is consistent with the observation that the ORF3-AAFIAA mutant retained partial palmitoylation and could partially rescue HEV progeny release (Fig. 4).

We acknowledge, however, that although our experimental data and modeling identify the ZDHHC17 ARD as the primary determinant of ORF3 binding, a potential contribution of the C-terminal ZDHHC17 PDZ domain cannot be excluded. Future studies should therefore examine whether this domain further recruits or stabilizes ORF3 at the appropriate membrane environment, or supports its productive incorporation into a larger trafficking complex.

Many questions remain regarding the mechanisms governing vectorial HEV release. Here, we identify ORF3 palmitoylation by ZDHHC17 as a central molecular mechanism linking viral protein–host enzyme interactions to apical trafficking and directional HEV release, thereby highlighting novel potential targets for antiviral intervention. More broadly, this work establishes a framework for studying protein interactions involving disordered proteins. In addition, an automated image analysis pipeline for quantification of apical signals was implemented which will facilitate future insights into the vectorial trafficking machinery of polarized epithelial cells.

## Material & Methods

### Standard cell culture

The human hepatoma cell line S10-3 (a kind gift from Suzanne Emerson) and HEK293T (ATCC CRL-3216) were cultured in Dulbecco’s Modified Eagle Medium (DMEM, Gibco) + GlutaMAX-I supplemented with 10% heat inactivated (HI) fetal bovine serum (FBS, capricorn), here referred to as complete DMEM (cDMEM). HepG2/C3A cells (ATCC HB-8065) were cultured on collagen-coated cell culture vessels in cDMEM. Caco-2 (ATCC HTB-37) cells were cultured on collage-coated cell culture vessels in DMEM + GlutamMAX-I supplemented with 20% HI FBS. Polarized Caco-2 cells were cultured on collagen-coated cell culture inserts (1 µm pore, Greiner) until a TEER of > 1000 Ω×cm² was reached. HepaRG cells were cultured and differentiated as previously described^57^. Cell lines were validated by phenotypic screening and confirmed to be mycoplasma-free using a PCR detection kit (Abcam). All cells were maintained at 37°C in 95% humidity and 5% CO_2_ atmosphere.

### Generation of cell lines

Various ORF3 derivatives were cloned into the lentiviral expression plasmid pLVX (Takara Bio Europe) and shZDHHC17 was cloned into the dox-inducible plasmid pTRIPZ (Dharmacon) by site-directed mutagenesis. Mutations were introduced by an overlap extension PCR using the Phusion polymerase (New England Biolabs) based on the primers listed in Table 1. Lentiviruses were produced by transfecting HEK293T cells with plasmids encoding VSV-G, HIV gag/pol proteins, and the transgene using the JetPRIME reagent (Polyplus) according to the manufacturer’s protocol. Lentiviruses were harvested 48 h post-transfection. Caco-2 cells were transduced with pLVX-ORF3 (WT or derivative) or pTRIPZ-shZDHHC17 and selected in cDMEM supplemented with 20% HI FBS and puromycin (10 µg/mL). Transduced cells were treated with 2 µg/ml dox (Sigma) for 48 h to induce shZDHHC17 expression.

**Table 1:**
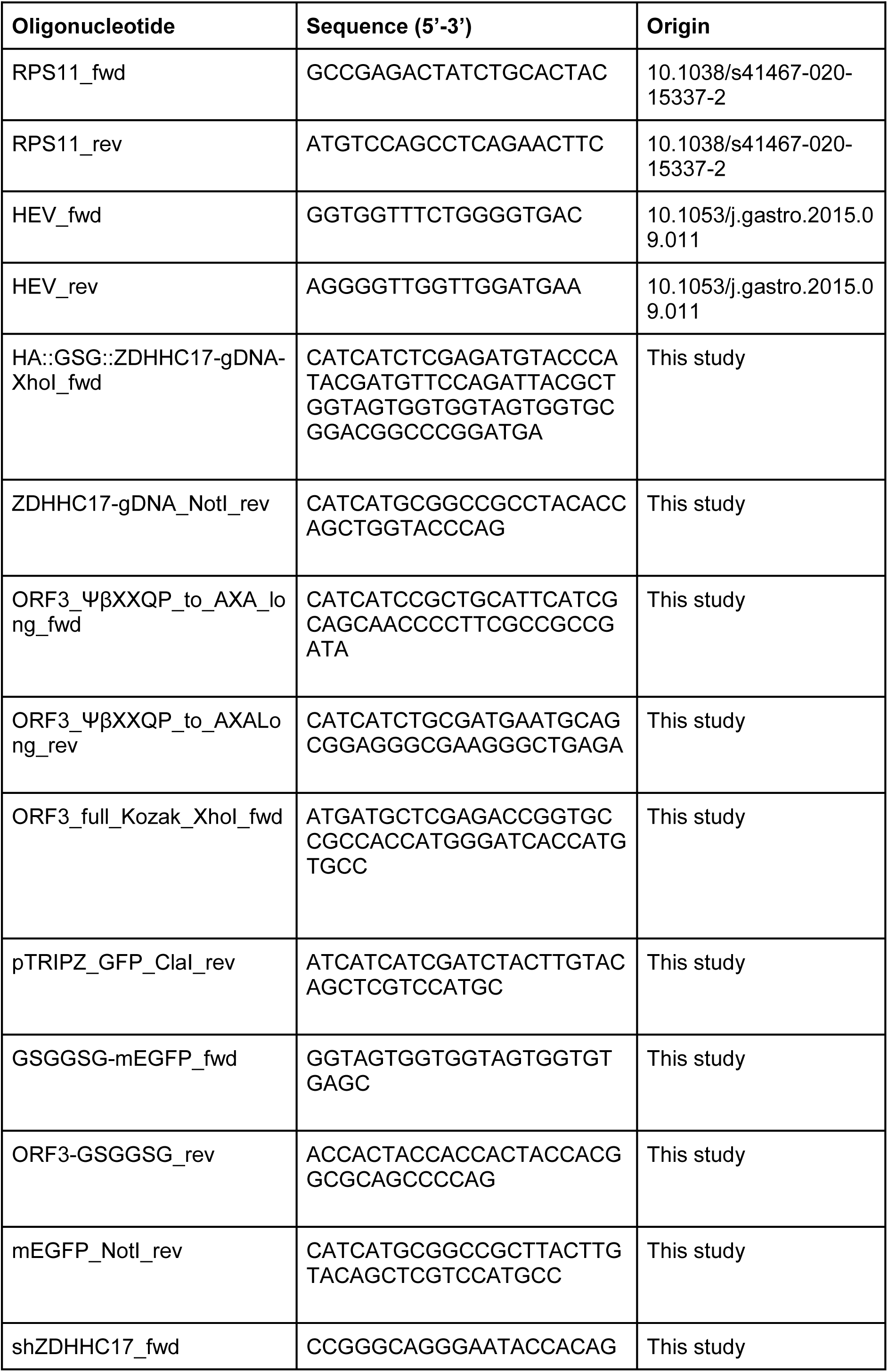
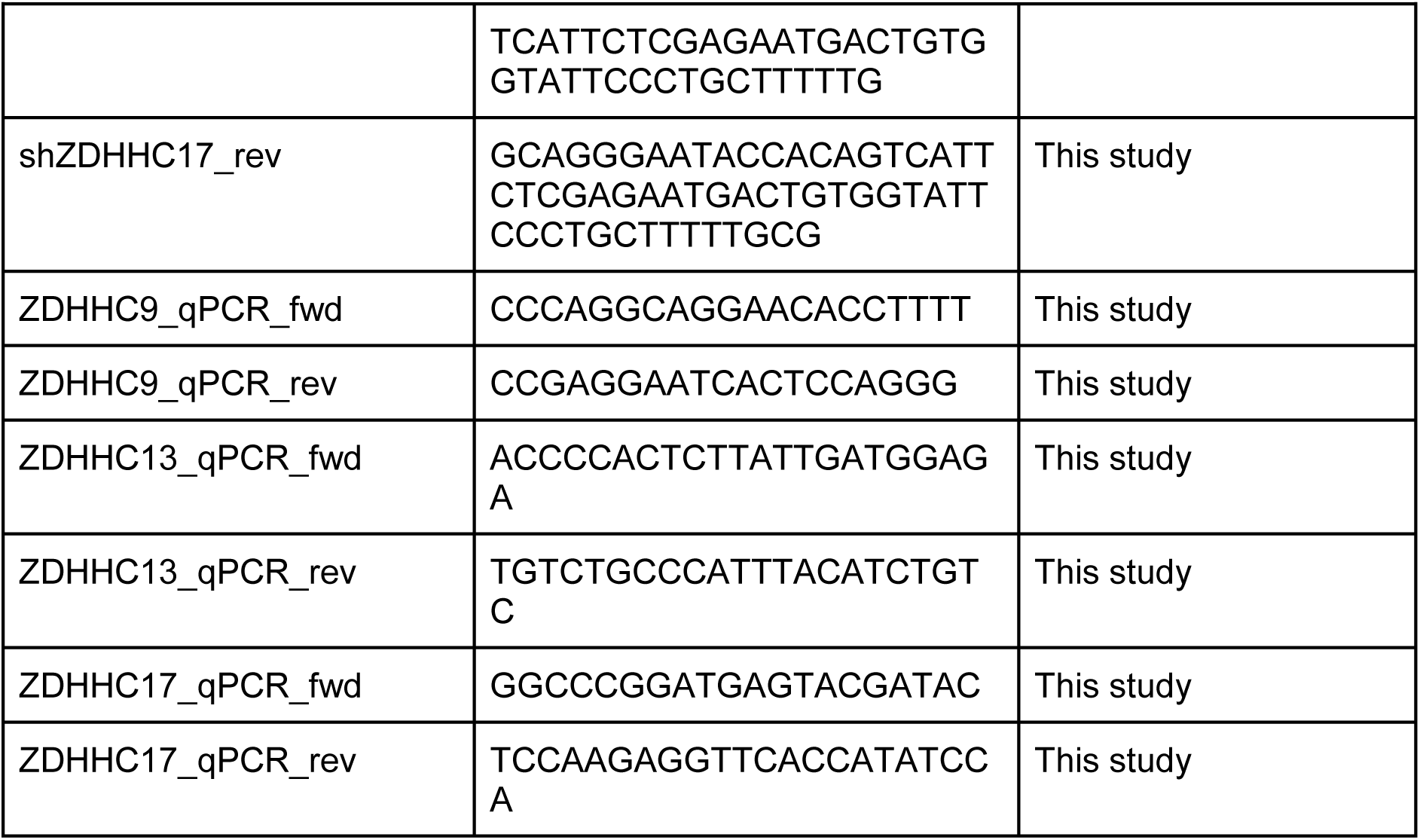
List of oligonucleotides used in this study.

### Immunofluorescence staining

Cells were fixed with 4% paraformaldehyde (PFA, Science Services) for 15 min at room temperature (RT). Cells were blocked and permeabilized with PBTG (10% goat serum, 1% bovine serum, and 0.1% TritonX-100 in PBS) for 30 min, followed by the incubation with primary antibody diluted in PBTG at 4 °C over night. After three washes with PBS, cells were incubated with the respective Alexa-conjugated secondary antibody diluted in PBTG for 1 h at RT. Cells on coverslips were mounted using ProLong^TM^ Glass Antifade Mountant (Thermo Fisher Scientific) and cured for at least 12 h in the dark. Transwells were stained in a wet chamber. After staining, the membrane was cut and mounted onto a coverslide with a coverslip using SlowFade™ Glass Soft-Set Antifade Mountant. The coverslips were sealed using Nail polish or Twinsil (picodent ® Dental-Productions- und VertriebsGmbH) and cured overnight in the dark.

### Image acquisition

For polarized cells and non-polarized cells, multichannel z-series with a spacing of at least 0.13 µm were acquired on a Zeiss Airyscan 2 LSM900 confocal microscope. A 63x oil immersion objective was used for all images. To determine foci focus units (FFUs) transparent 48 well-plate (Greiner Bio-One) was imaged using a Zeiss Celldiscoverer 7 inverted light microscope. Images were analyzed and merged with Fiji^58^.

### Quantitative image analysis

3D image stacks were cropped to exclude regions below the cell, defined by the lowest z-slice with the highest mean MemBrite® or GFP signal (for some images with unusual intensity distribution this was performed manually). The images were also cropped along the x- and y-axes by selecting a region of interest (ROI), which was performed by maximum-intensity projection of the GFP channel of the 3D image along the z-axis (excluding the lowest five z-slices), thresholding the resulting maximum-intensity projection image using Otsu’s method^36^, morphological hole filling, and connected component analysis. If the largest connected component covered 5–20% of the maximum-intensity projection image, the bounding box of this component was expanded symmetrically by 25% along the x-axis and the y-axis and then selected as ROI (otherwise, the image was not cropped along the x- and y-axes). The apical region was identified based on enriched MemBrite® signal (∼2.5 µm along the z-axis). For localizing the top border of the apical region, the MemBrite® channel was divided into 12×12 pixel patches in the xy-plane. Within each patch, mean MemBrite® intensities along the z-axis were calculated and the uppermost z-position satisfying one of two conditions was selected as the top border of the apical region: (1) The membrane intensity at this z-position and the next two z-positions below exceeded a threshold, or (2) the membrane intensity at this z-position and the next z-position below exceeded a threshold, and also the intensity at the next z-position below was higher than that at the next z-position above. The threshold was determined adaptively by using the mean value of the membrane intensity profile, or half of the mean if the intensity distribution was right-skewed. Outliers were corrected using median values from neighboring patches. The apical region was defined as a band below the localized top border, and the non-apical region as the region below it. To reduce image noise, GFP signals were clipped at 99.9^th^ percentile of the GFP intensity distribution, smoothed (Gaussian filter, σ = 1 pixel), and the background signal was segmented using Otsu thresholding^36^. The GFP intensities were then quantified within apical and non-apical regions, excluding background pixels, and the percentage of apical GFP over the total apical and non-apical signal was computed.

### Sample preparation for affinity purification of HEV-ORF3 proteins and cellular proteins

For HEV–ORF3 interactome analysis, four independent affinity–purification experiments were performed in HepG2/C3A cells transduced with lentivirus to express either pLVX–ORF3 or pLVX–mEGFP, each C–terminally HA–tagged. Cells were expanded in quadruplicate (two confluent 15–cm dishes per replicate), washed with ice–cold PBS, scraped, pooled by replicate, and pelleted at 500 × g for 5 min at 4 °C. Pellets were washed twice, pellets were lysed in buffer containing 0.2% NP–40, 100 mM NaCl, 5% glycerol, 1.5 mM MgCl₂, and 50 mM Tris–HCl (pH 7.5), with benzonase and EDTA–free Complete Protease Inhibitor (Roche), incubated 30 min on ice, sonicated (Bioruptor, high intensity, 30 s on/off for 5 min), and clarified at 15,000 × g for 30 min at 4 °C. Protein concentration was determined (Pierce 660 nm Assay, Thermo Fisher Scientific), and samples were adjusted to 2.5 mg total protein in 700 µL lysis buffer plus protease inhibitor. Cleared lysates were incubated overnight at 4 °C with 30 µL anti–HA agarose beads, followed by three washes with lysis buffer and three with wash buffer (100 mM NaCl, 5% glycerol, 1.5 mM MgCl₂, 50 mM Tris–HCl, pH 7.5). Bead–bound proteins were resuspended in 4% SDC buffer (4% sodium deoxycholate in 100 mM Tris–HCl, pH 8.5), heated at 95 °C for 5 min, reduced and alkylated with TCEP and CAA, and digested sequentially with LysC (3 h) and trypsin (15 h) at 37 °C. Peptides were desalted using SDB–RPS StageTips, eluted with 1.25% NH₄OH in 60% ACN, dried in a SpeedVac, resuspended in 0.1% FA, and quantified at 280 nm before LC–MS/MS.

### Liquid chromatography-tandem mass spectrometry (LC–MS/MS)

LC–MS/MS analysis was performed on an EASY-nLC 1200 system (Thermo Fisher Scientific) coupled online to a Q Exactive HF–X mass spectrometer via a nano–electrospray source^59^. Peptides were separated on a 20–cm C18–AQ analytical column (1.9 µm, 75 µm i.d.; Dr. Maisch) at 300 nL/min, using 0.1% formic acid in water (A) and 0.1% formic acid in 80% ACN (B). A gradient of 5–30% B over 85 min, 30–60% over 12 min, 60–80% over 3 min, and 80–95% over 1 min was followed by a 5–min hold at 95% B, with re–equilibration to 5% B. After each run the column was washed for 15 min with 95% B to minimize carryover. Data were acquired in data–dependent mode (DDA) over m/z 300–1,650 at 60,000 resolution (AGC 3 × 10⁶); the 15 most intense precursors were fragmented by HCD (NCE 27%) and MS/MS spectra recorded at 15,000 resolution (AGC 1 × 10⁵, max 25 ms, dynamic exclusion 20 s).

### Data processing and statistical analysis of MS samples

AP–LC–MS/MS data were processed using MaxQuant (v2.6.3.0) with the built–in Andromeda search engine as previously described^60^. Label–free quantification (LFQ) and “match between runs” were enabled, and MS/MS spectra were searched against a database comprising forward and reverse sequences of the human proteome (UniProtKB UP000005640, Sep 2023), HEV proteins (UniProt P29324 and related entries, Apr 2024), mEGFP, and the HA tag. Carbamidomethylation of cysteine was set as a fixed modification; oxidation of methionine and N–terminal acetylation were set as variable modifications. Precursor and fragment mass tolerances were 6 ppm and 0.5 Da, respectively, and peptide–spectrum matches and protein identifications were filtered at 1% FDR. ProteinGroups output tables were analyzed in Perseus (v2.0.11): reverse hits, potential contaminants, and proteins identified “only by site” were removed; LFQ intensities were log₂–transformed, and only proteins quantified in at least three replicates in at least one group were retained. Missing values were imputed from a normal distribution (width 0.3, downshift 1.8). Differential enrichment between HEV–ORF3 and mEGFP pulldowns was evaluated using a two–sided Student’s t–test with permutation–based FDR correction, and proteins significantly enriched in HEV ORF3 samples were considered candidate interacting partners. The volcano plot was generated in R.

### Co-immunoprecipitation

1.2 x 10^7^ HEK293T cells were seeded on 10-cm cell culture dishes coated with 100 µg/mL poly-L-lysine (Sigma-Aldrich). The next day, cells were co-transfected with 10 µg of pLVX encoding ORF3::GFP or GFP-tagged ORF3 derivatives and 10 µg of the pLVX encoding HA-ZDHHC17 in cDMEM without pen/strep. JetPrime (Polyplus ®) was used for transfection at a DNA:JetPrime ratio of 1:1. Medium was changed after 5–6 h. After 24 h, the cells were lysed in ice-cold lysis buffer (10 mM Tris/Cl pH 7.5, 150 mM NaCl, 0.5 mM EDTA, 0.5% Nonidet™ P40 Substitute), supplemented with 1x cOmplete Mini Protease Inhibitor Cocktail (Roche) and incubated for 30 minutes on ice, agitated every 10 minutes. After removal of DNA, 10% of lysate was set aside as input and boiled with Laemmli sodium dodecyl sulfate (SDS) sample buffer at 95 °C for 10 min. 15 µL GFP-Trap magnetic agarose (proteintech) were added to each sample and incubated for 2 h, at 4 °C. Beads were washed five times with wash buffer (10 mM Tris/Cl pH 7.5, 2 M NaCl, 2 % Nonidet™ P40 Substitute, 1% SDS, 10% Glycerol, 0.5 mM EDTA) and eluted in Laemmli SDS sample buffer by boiling at 95 °C for 5 min.

### AlphaFold modeling

Full-length models with and without palmitoylation were generated using AlphaFold3 with default parameters (5 models per run)^61^. Membrane insertion was predicted using the PPM 3.0 web server, employing a predefined membrane composition representative of the Golgi apparatus. To further improve models, AlphaFold2 predictions were performed using ColabFold (v1.5.5) through the MassiveFold implementation, allowing the use of different model parameter sets on a truncated version of ORF3 (residues 37–83) and the ARD of ZDHHC17 (residues 55–288)^62,63^. Multiple sequence alignments were generated with MMseqs2 using the UniRef30 database (release 2302) and the environmental database (release 2021_08)^63–65^. All analyses were carried out using in-house Python scripts, and molecular graphics were generated with PyMOL^66^.

### Molecular dynamic simulation

Initial ORF3 complex structures were selected from MassiveFold predictions based on pLDDT and conformation (models 0, 1, 10, and 16). Systems were parameterized with the Amber ff99SB-ILDN force field and simulated using GROMACS 2025^67^.Each system was solvated in a triclinic box (22 Å padding) with TIP3P water and 0.15 M NaCl^68^. After two-step energy minimization (restrained and unrestrained), equilibration was performed with 100 ps NVT and 100 ps NPT at 300 K using V-rescale and C-rescale, respectively^69,70^. For each model, three independent 250 ns production runs were carried out (10 ps output). Electrostatics were treated using the zero-dipole method (12 Å cutoff), with a 4 fs time step enabled by hydrogen mass repartitioning. LINCS and SETTLE were applied, and the center of mass was restrained as described previously^71^.System details are listed in Sup. Table 1.

## Data availability

Proteomic data sets have been deposited to PRIDE with project accession: PXD078645. Token: (only accessible to reviewers). Alternatively, reviewer can access the dataset by logging in to the PRIDE website using the following account details: (Only accessible to reviewers). Source data are provided with this paper. All scripts used AlphaFold structural models and molecular dynamics simulations data produced in this study with are available on BSM-Arc (only accessible for reviewers))

## Acknowledgements

The authors gratefully acknowledge Suzanne Emerson, Jérôme Gouttenoire, Darius Moradpour, Hans-Georg Kräusslich and Rainer Ullrich for sharing reagents. The authors acknowledge the help of Carola Krug for contributing to the development of the quantitative imaging analysis pipeline. The authors acknowledge the data storage service SDS@hd supported by the Ministry of Science, Research and the Arts Baden-Württemberg (MWK) and the German Research Foundation (DFG) through grant INST 35/1803-1 FUGG and INST 35/1804-1 LAGG. This project was supported by GENCI at CINES and Idris with HPC and storage resources, thanks to the grant AD010714400R1 on the supercomputer (Jean Zay and Adastra’s A100 and MI250x partition). This work was in part performed under the International Collaborative Research Program of the Institute for Protein Research, Osaka University, ICR-25-21. Computational resources from the TSUBAME4.0 system (Institute of Scientific and Industrial Research, Tokyo) were provided by the HPCI Research Project hp250165. We acknowledge Vibor Laketa, head of the Infectious Disease Imaging Platform (IDIP) at the University Hospital for external support.

## Funding

This work was supported by grants from the Deutsche Forschungsgemeinschaft (DFG, German Research Foundation) - Projektnummer - 240245660—SFB 1129 (to C.F., K.R, and V.L.D.T) and Projektnummer - 272983813-TRR 179 (A.P. and V.L.D.T.). V.L.D.T. was further supported by the Chica and Heinz Schaller Foundation.

## Author contributions

Conceptualization, P.B., C.C.S., V.L.D.T.; Methodology, P.B., C.C.S., C.F., K.R., V.L.D.T.; Investigation, P.B., C.S., D.G., A.Piras., Z.E., A.F., H.C., P.J., C.F., S.P.T., T.T.; Resources, V.L.D.T., A. Pichlmair; Software, C.K., L.K., S.P.T., T.T.; Formal analysis, P.B., C.S., C.K., L.K., D.G., A.Piras., C.F., S.P.T., T.T.; Writing-original draft, P.B., V.L.D.T., T.T.; Final draft, P.B., V.L.D.T., Supervision, K.R, A.P., V.L.D.T.; Funding, V.L.D.T.

## Competing interests

Authors declare that they have no competing interests.

## Notes

### Competing Interest Statement

The authors have declared no competing interest.

### Summary of Updates

Informations for reviewers have been adapted.

