## Supplementary data for "ZDHHC17–Mediated Palmitoylation of Hepatitis E Virus ORF3 Protein Regulates Vectorial Trafficking in Polarized Epithelial Cells"

### Supplementary Methods

*Generation of human induced pluripotent stem cell-derived HLCs and intestinal organoids*

The human induced pluripotent stem cell (iPSC) line PSC.3A (a gift from S. Duncan) was maintained in mTeSR1 medium (STEMCELL Technologies) on plates coated with Matrigel (Corning) according to the manufacturer's instructions. Cells were differentiated into small intestinal organoids[1] or hepatocyte-like cells (HLCs[2]) as described previously[3]. This work with human iPSCs (S-439/2018) was approved by the Ethics Committee of the Medical Faculty of Heidelberg University.

*Western blot*

Caco-2 were lysed in RIPA lysis buffer (Thermo Fisher) supplemented with 1x cOmplete Mini Protease Inhibitor Cocktail (Roche) on ice for 30 minutes, supplemented with Laemmli SDS buffer, and boilder at 95°C for 10 min and loaded onto a SDS-PAGE gel. The proteins were transferred to 0.2 µm pore size polyvinylidene difluoride (PVDF, G-Biosciences) membranes by wet blotting using standard methods. The membranes were blocked with 5% milk/0.1% Tween-20 in PBS (PBS-T). The following antibodies were incubated overnight in 5% milk/PBS-T at 4°C: mouse α-ORF3 1:100 (University of Geneva Antibody Facility, cat. no. ABCD_RB198); mouse α-actin (Sigma) 1:4000; mouse α-HA (Proteintech) 1:4000, rabbit α-GFP (a kind gift from Hans-Georg Kräusslich) followed by staining with corresponding secondary antibodies conjugated with HRP (Jackson ImmunoResearch). Membranes were imaged with Pierce™ Enhanced Chemiluminescence (ECL) Western blotting substrate and images were acquired using ChemoStar Touch ECL & Fluorescence Imager (Intas).

*High-pressure freezing and immunogold electron microscopy*

Cells were seeded onto UV-sterilized carbon-coated sapphire discs (3 mm diameter, 0.05 mm thickness) at a density of $6.3\times10^4$ cells/cm$^2$. After 2 days, each sapphire disc was sandwiched between two aluminum carriers and subjected to high-pressure freezing using an HPM 010 system, with the 0.1 mm deep carrier placed on the cell side and the flat 0.3 mm carrier positioned beneath the sapphire. High-pressure freezing and the subsequent immunogold electron microscopy[4]. Indirect immunogold labeling was performed on 70 nm sections using a non-commercial anti-GFP primary antibody (GFP-Q, gift from the Center for Organismal Studies, Heidelberg). Sections were fixed with glutaraldehyde, post-stained with uranyl acetate and lead citrate, and examined using a JEOL JEM-1400 transmission electron microscope operated at 80 kV and equipped with a TemCam F416 4K camera.

*In vitro transcription and EPO of HEV WT and ΔORF3 RNA*

pBSK-HEV-p6-derived plasmids were linearized with MluI, respectively, and RNA was in vitro transcribed using the mMESSAGE mMACHINE T7 kit (Invitrogen). 4 x $10^6$ Caco-2 cells were electroporated at 270 V and 975 µF with 5 µg of IVT HEV RNA in Cytomix (120 mM KCl, 0.15 mM CaCl2, 10 mM KPO4, 25 mM HEPES, 2 mM EGTA, and 5 mM MgCl2), supplemented with adenosine triphosphate (ATP) and glutathione (GT). Cells were resuspended in cDMEM and cultured for 5 days until seeded in a 24-well plate for siRNA treatment as described above.

*Quantitative (real time) reverse transcription PCR (RT-qPCR)*

RNA was extracted from cell lysates with the Universal RNA kit (Roboklon) following respective manufacturer's instructions. cDNA was synthesized using the iScript cDNA Synthesis Kit (Bio-Rad) or the High Capacity cDNA Reverse Transcription Kit (Thermo Fisher Scientific) and diluted. qPCR was then performed with iTaq Universal SYBR Green Supermix (Bio-Rad) on a CFX96 Real-Time PCR Detection System (Bio-Rad) using the primers listed in Table 1. Absolute HEV genome copies were calculated from an HEV standard curve, produced by serial 10-fold dilutions of a 10 ng/µL-concentrated pBSK-HEV-p6 plasmid. ZDHHC9, ZDHHC13 and ZDHHC17 expression in Caco-2 cells were normalized over the housekeeping gene RPS11 using the $2^{-\Delta Ct}$ method and additionally normalized to siNT using the $2^{-\Delta\Delta Ct}$ method.

*Virus production & Infection*

Production of cell culture HEV-3 Kernow-C1/p6 strain virus was conducted as previously described[1]. Cells were infected with a MOI of 1, calculated from FFU/mL.

*shRNA/siRNA treatment and titration*

SMARTpool siRNAs (Dharmacon) were individually added to each collagen-coated well of a 24-well plate containing 48 µL OptiMEM (Gibco) and 2 µL Lipofectamine™ RNAiMAX Transfection Reagent (Thermo Fisher) at a final concentration of 100 nM. After 5 min of incubation of room temperature, Caco-2 cells ($7.5x10^4$ in 300 µL of complete DMEM (cDMEM)), were added to each well. After 48 h of incubation a media change was conducted and 500 µL of fresh DMEM (supplemented with 10 % FBS, 1 % Penicillin and Streptomycin) was added. 96 h post-transduction the supernatant was collected in an Eppendorf tube and titrated on HepG2/C3A cells. 5 days post-infection the cells were washed twice with PBS. The cells were harvested by scraping in 200 µL of PBS per well. Using an inducible shRNA, the p6 RNA electroporated cells were induced on day 0 with Doxycyclin (DOX) (1 µg/mL) for 48 h. The media was exchanged and 48 h later the cells and supernatant were harvested.

**Supplementary Figures**

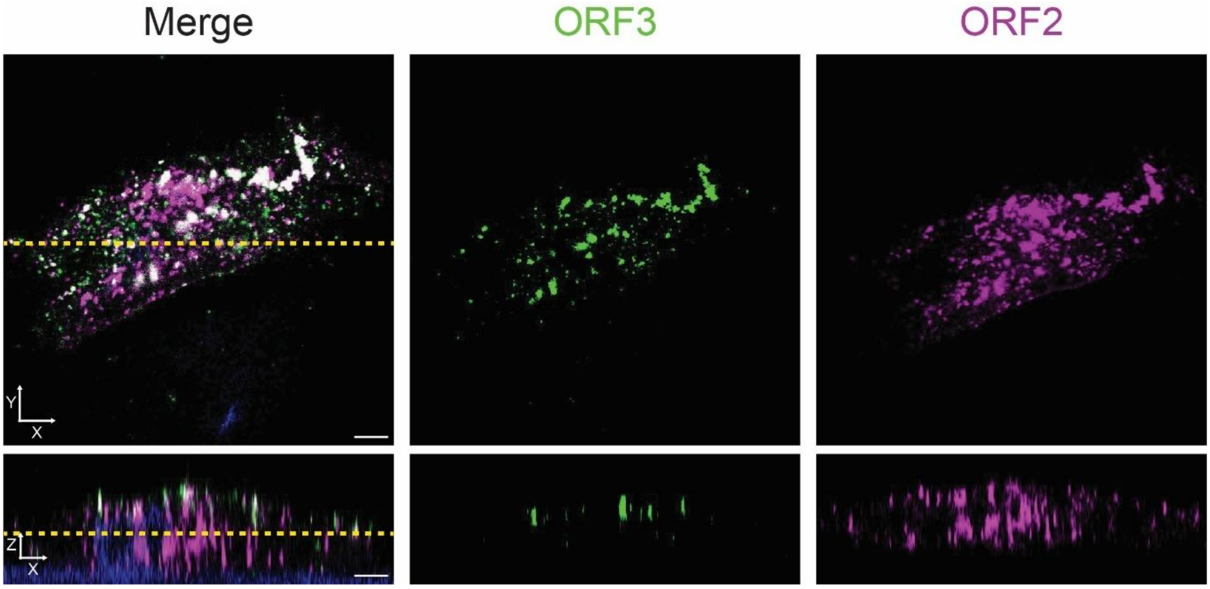


**Supplemental Figure 1. ORF3 localizes to the apical membrane in hepatocyte-like cells (HLC)**. HEV-infected HLCs were fixed 7 days post-infection and stained for ZO1 (yellow), ORF3 (green), ORF2 (magenta) and nuclei (blue). Images were taken with an Airyscan 2 LSM 900. Shown are single slices and XZ view (indicated by the yellow line). Scale bar = 5 µm.

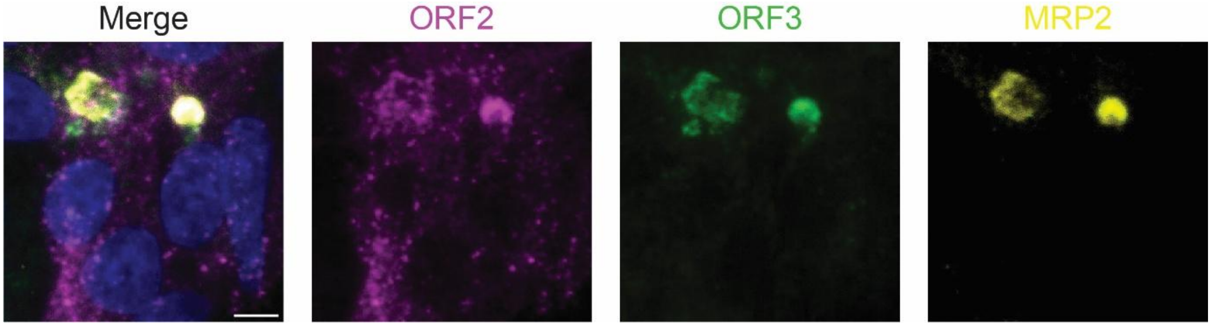


**Supplemental Figure 2. ORF3 co-localizes with MRP2 in differentiated HepaRG.** HEV-infected HepaRG were fixed 7 days post-infection and stained for MRP2 (yellow), ORF3 (green), ORF2 (magenta) and nuclei (blue). Images were taken with an Airyscan 2 LSM 900, single slices are shown. Scale bar = 5 µm.

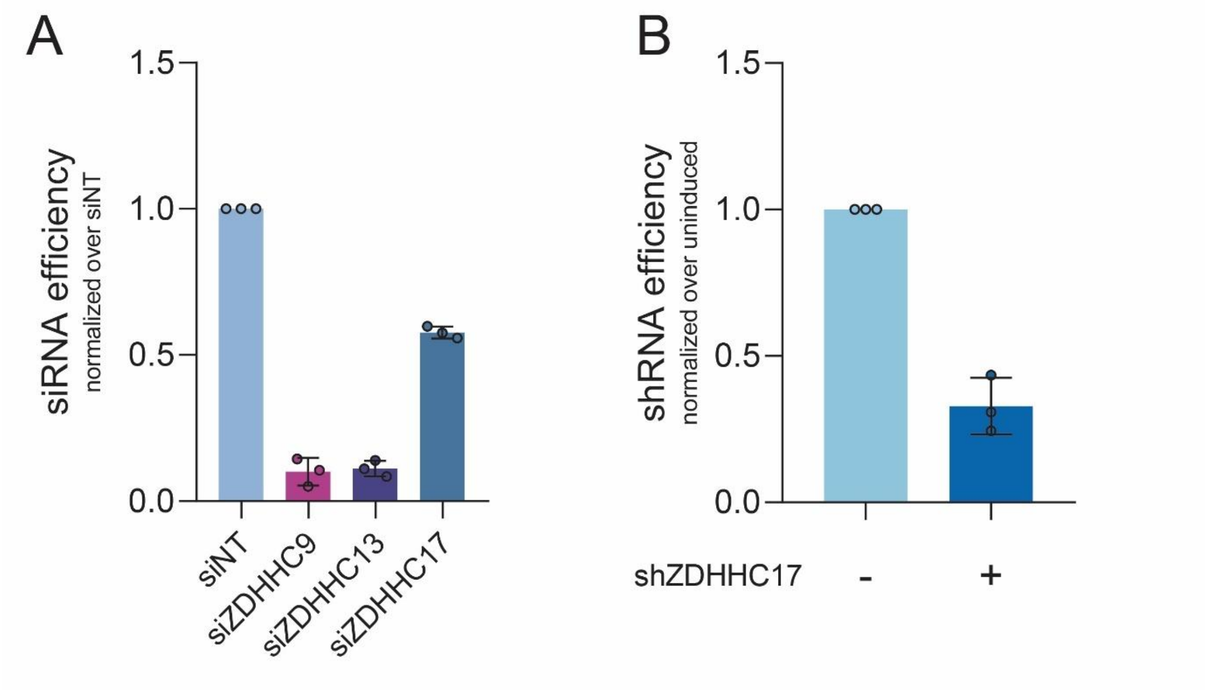


**Supplemental Figure 3. Knockdown efficiency of ZDHHC transferases.** Knockdown efficiency in Caco-2 cells of **(A)** siZDHHC9, siZDHHC13, siZDHHC17 and **(B)** shZDHHC17 was assessed by RT-qPCR using the primers in Table 1. Cells were (A) transfected with siRNA or (B) induced with dox for 2 days, media was replaced, and cells were harvested after an additional 2 days.

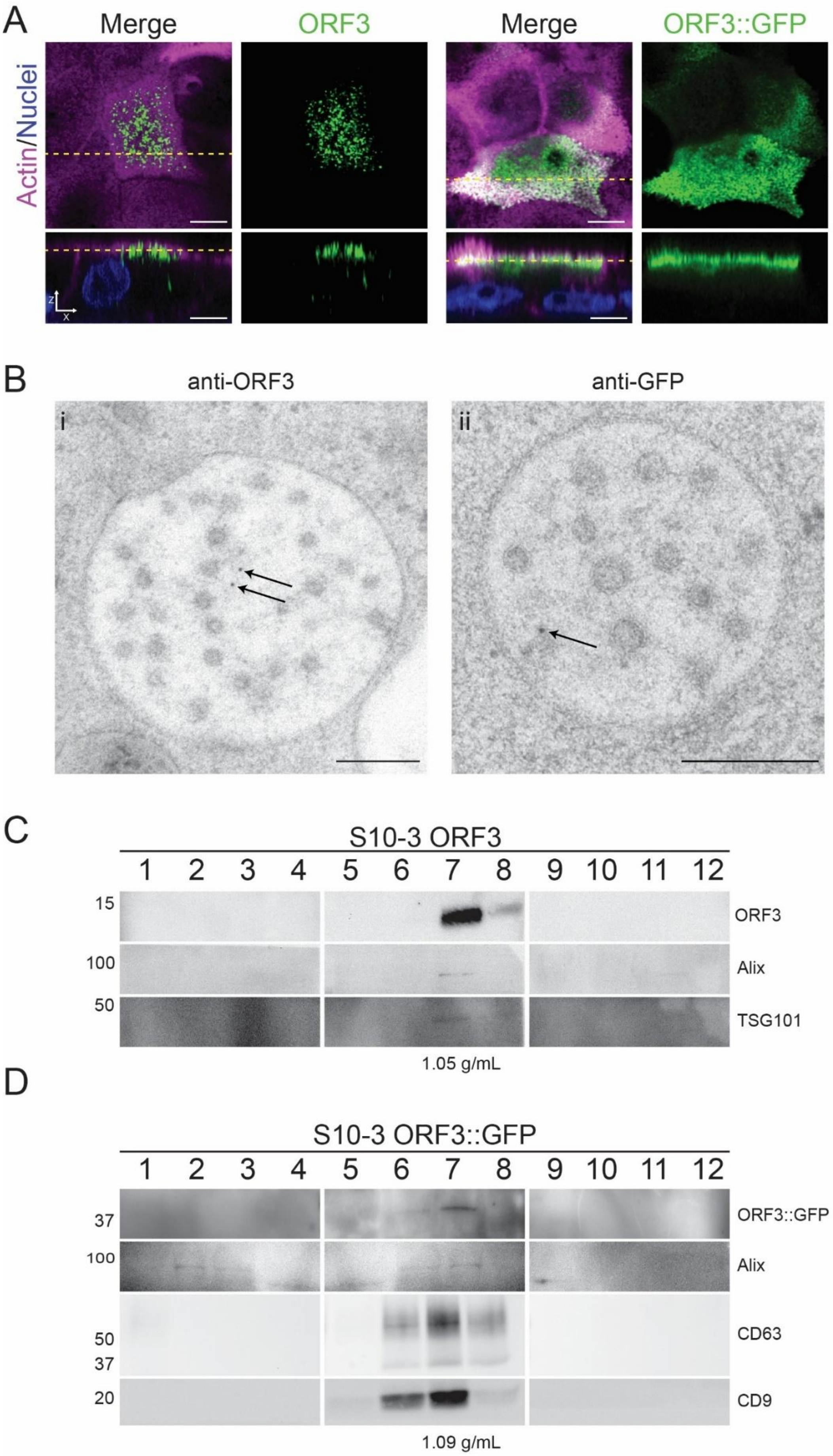
A
Merge
ORF3
Merge
ORF3::GFP
Actin/Nuclei
B
anti-ORF3
anti-GFP
i
ii
C
S10-3 ORF3
1 2 3 4 5 6 7 8 9 10 11 12
15
100
50
ORF3
Alix
TSG101
1.05 g/mL
D
S10-3 ORF3::GFP
1 2 3 4 5 6 7 8 9 10 11 12
37
100
50
37
20
ORF3::GFP
Alix
CD63
CD9
1.09 g/mL

**Supplemental Figure 4. GFP-tagging does not impair ORF3 apical localization or secretion via multivesicular bodies. (A)** Caco-2 cells stably expressing ORF3 or ORF3::GFP and fixed after polarization (TEER of > 1000 Ω x $cm^2$) the cells were fixed, stained for actin and nuclei. Super resolution images were taken with a Zeiss Airyscan 2 LSM 900. Green: ORF3 or ORF3::GFP, Magenta: Actin, Blue: Nuclei. Dotted lines indicate XZ view. Scale bar = 5 µm. **(B)** EM images of MVBs of S10-3 cells stably expressing ORF3::GFP with anti-ORF3 (i) or anti-GFP (ii) primary and colloidal gold-coupled secondary antibody (black dots). Scale bar = 200 nm. **(C, D)** Supernatants of S10-3 cells stably expressing ORF3 (C) or ORF3::GFP (D) were differentially centrifuged and the extracellular vesicle-containing pellet was fractionated on an iodixanol gradient. The fractions were analyzed by western blot with antibodies against ORF3, GFP, Alix, TSG101, CD63 and CD9.

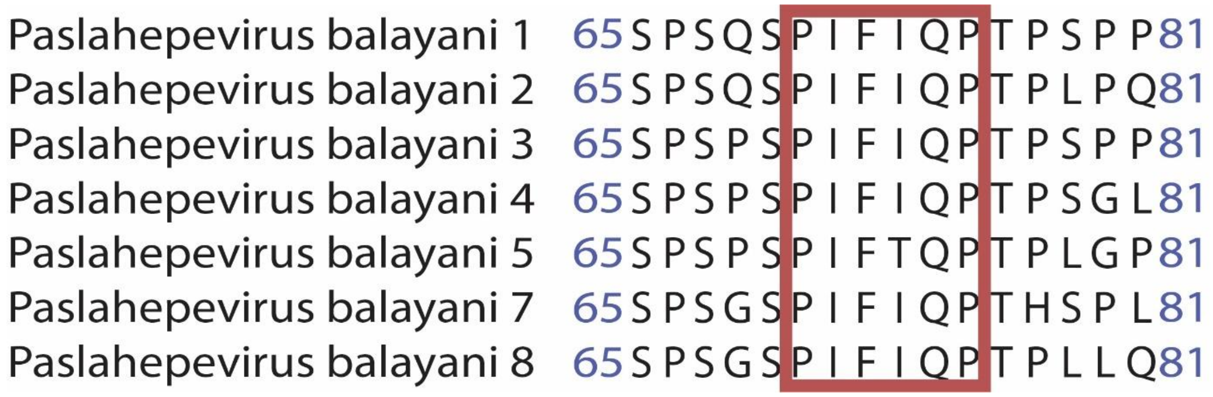


**Supplemental Figure 5. Conservation of ΨβXXQP among ORF3.** Sequence alignment using Jalview (Clustal Omega) of ORF3 from different strains of the genus *Paslahepevirus balayani*. Motif is indicated by red square.

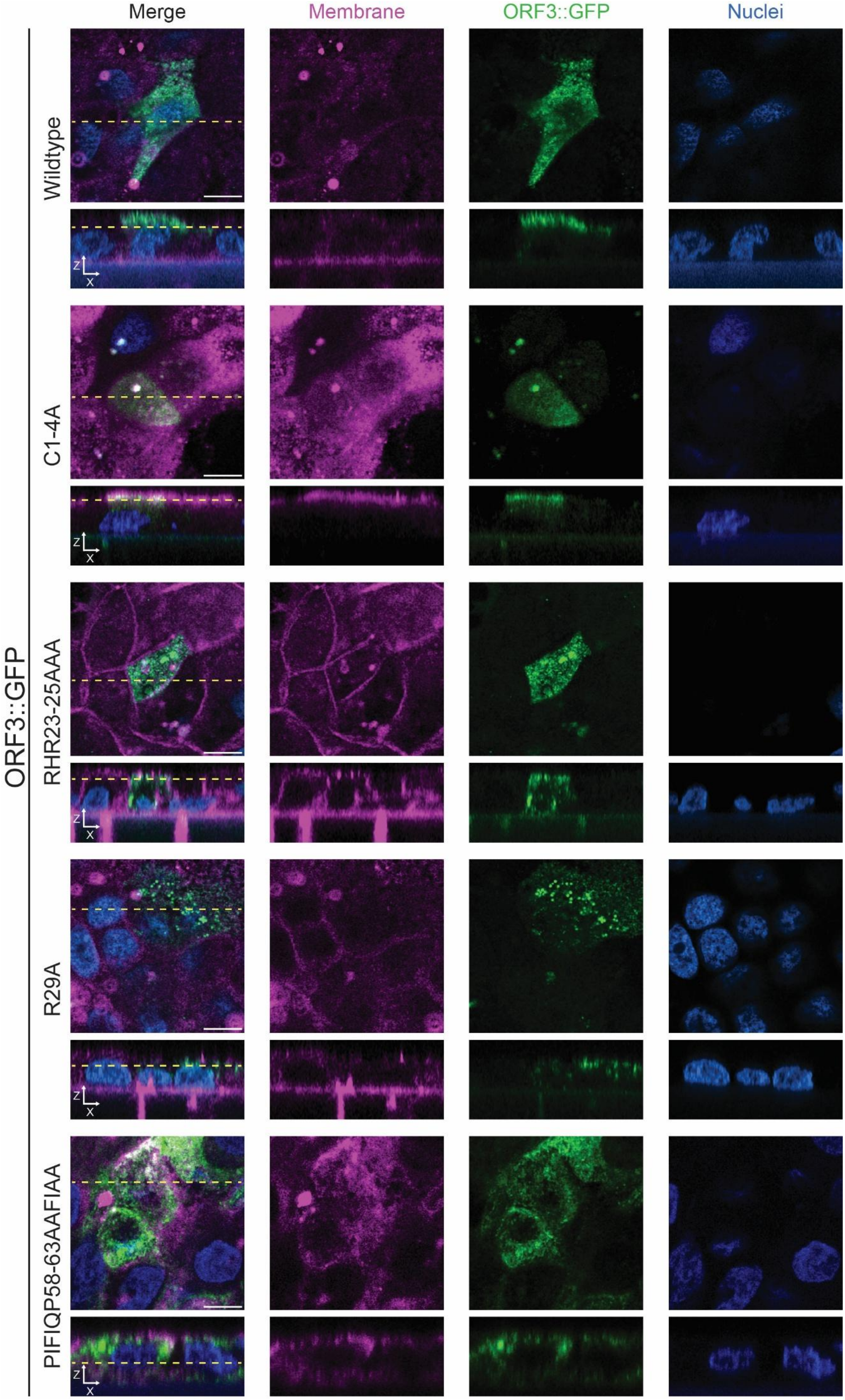
Merge
Membrane
ORF3::GFP
Nuclei
ORF3::GFP
Wildtype
C1-4A
RHR23-25AAA
R29A
PIFIQP58-63AAFIAA
z
x

**Supplemental Figure 6. Representative images of Caco-2 cells expressing ORF3::GFP variants used in the image-analysis pipeline.**

Z-stack images used for quantitative analysis of apical ORF3::GFP localization in polarized cells. Membrite®Fix staining (magenta) was used to define the apical membrane region, ORF3::GFP variants are shown in green, and nuclei are shown in blue. The pipeline quantifies the proportion of ORF3::GFP signal in the apical region relative to total cellular signal. Scale bar = 5 µm.

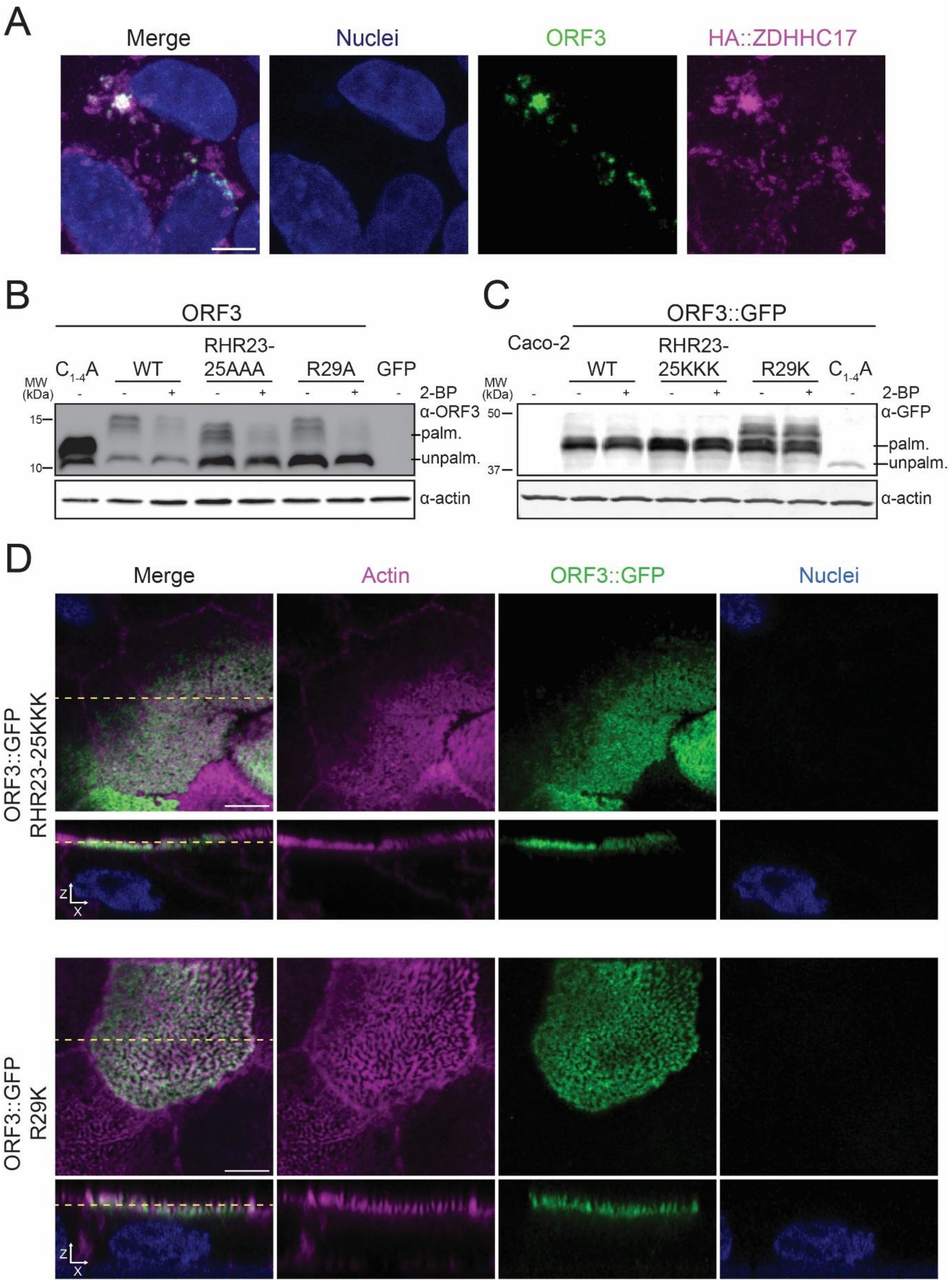


**Supplemental Figure 7. ORF3 colocalizes with ZDHHC17 and positively charged amino acids are important for ORF3 palmitoylation. (A)** Non-polarized Caco-2 cells constitutively expressing HA-tagged ZDHHC17 were electroporated with HEV RNA, fixed at 7 days post-infection and stained for HA (magenta), ORF3 (green), and nuclei (blue). Scale = 5 µm. **(B)** HEK293T cells were transfected to express ORF3 WT or derivatives. After 5 h, cells were treated with the palmitoylation inhibitor 2-bromopalmitate (2-BP) or DMSO and analyzed by

Western blot 24 h post-transfection. GFP-transfected cells served as negative control. **(C)** Caco-2 cells stably expressing DOX-inducible ORF3::GFP WT or derivatives were induced with DOX and treated with 50 µM 2-BP or DMSO. After 24 h, protein expression was analyzed by Western blot. Arrows indicate palmitoylated (upper bands) and non-palmitoylated (lower band) ORF3 species. Caco-2 WT cells served as negative control. (D) Confocal image of polarized Caco-2 cells expressing dox-inducible ORF3::GFP variants (RHR23-25KKK and R29K) were dox-induced after reaching a TEER of> 1000 Ω x $cm^2$. After 24h cells were fixed and stained for actin, and imaged using a Zeiss Airyscan 2 LSM 900 microscope. Scale bar = 5 µm.

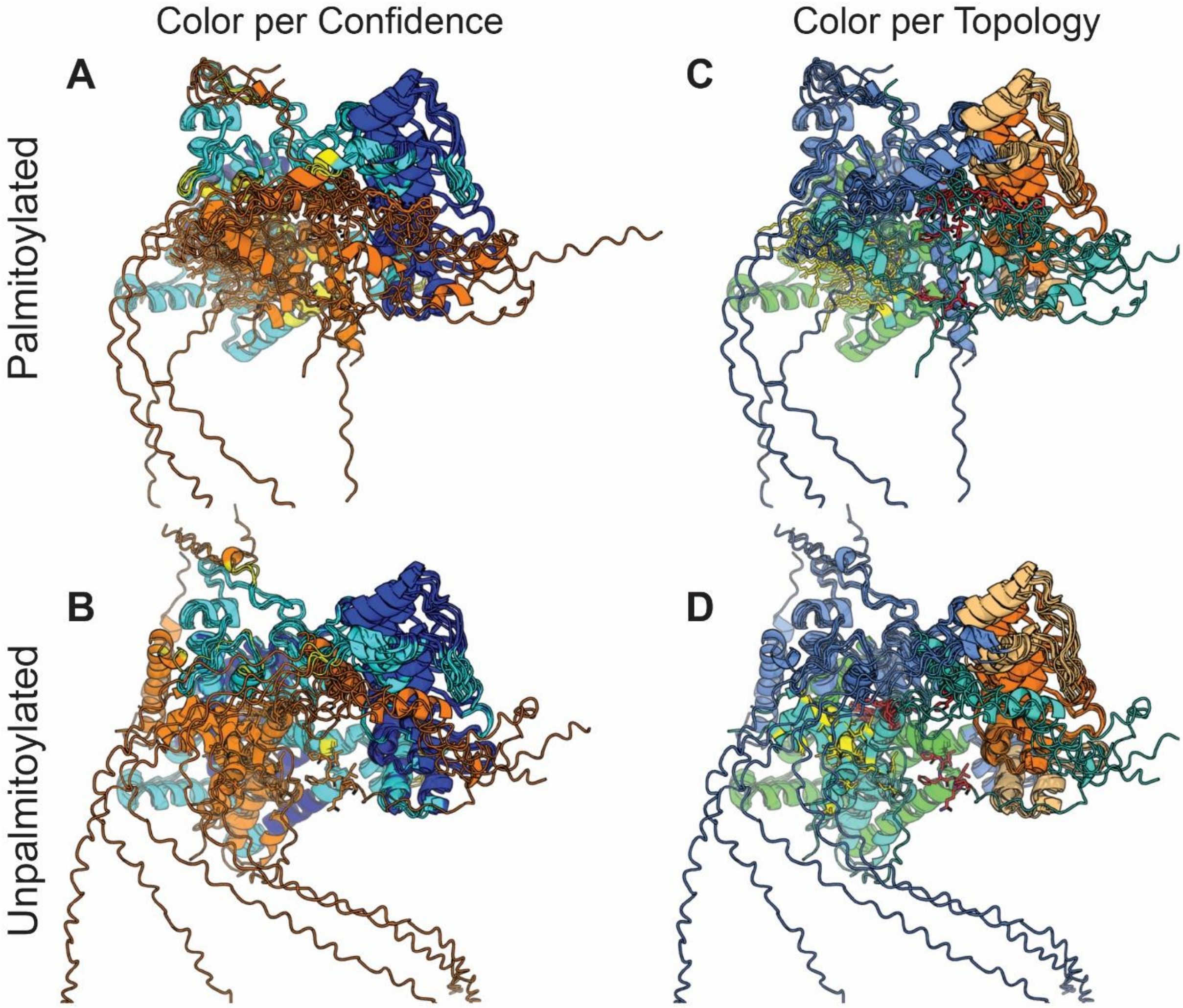


**Supplemental Figure 8. Comparison of palmitoylated and unpalmitoylated ORF3–ZDHHC17 models.** AlphaFold3 models of the ORF3–ZDHHC17 complex are shown in palmitoylated (top, **A**, **C**) and unpalmitoylated (bottom **B**, **D**) conditions. Left panels display models colored by AlphaFold confidence (pLDDT, A, B): very high confidence (pLDDT > 90) in dark blue, high confidence (70–90) in light blue, low confidence (50–70) in yellow, and very low confidence (< 50) in orange. Right panels show the same structures colored by topology (C, D): ZDHHC17 is shown in blue, with its ankyrin repeat domain (ARD) highlighted in orange and its transmembrane domain in green. ORF3 is shown in turquoise, with the PIFIQP motif highlighted in red and palmitoylated cysteines in yellow.

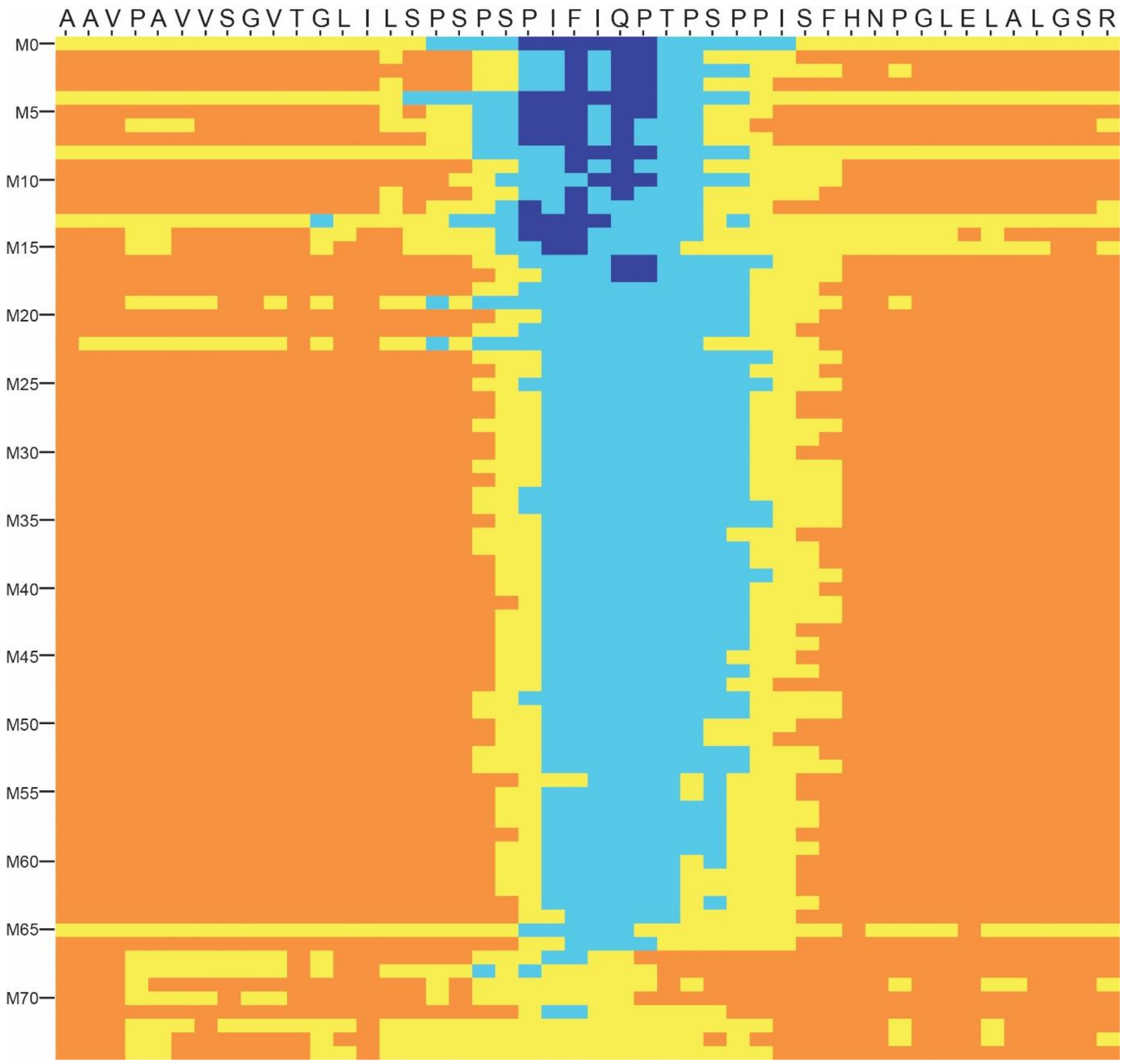


**Supplemental Figure 9. Heatmap of pLDDT scores across the top 75 models, highlighting high confidence (>90) in the PIFIQP region**. Extensive AlphaFold2 sampling of truncated ORF3 residues (residues 37-83). pLDDT confidence: very high confidence (pLDDT > 90) in dark blue, high confidence (70–90) in light blue, low confidence (50–70) in yellow, and very low confidence (< 50) in orange.

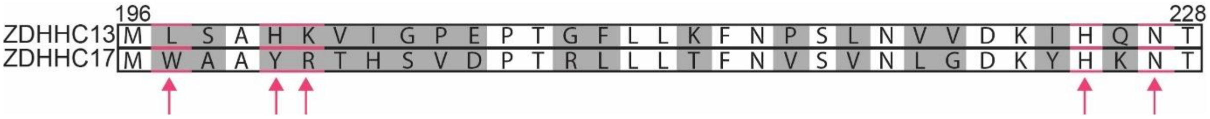


**Supplemental Figure 10. Alignment of ankyrin repeat domain 5 of ZDHHC13 and ZDHHC17**. Sequence alignment generated using Geneious Prime. Amino acids differing between both proteins are highlighted in grey. Residues identified by molecular dynamics as important for the interaction between ORF3 and ZDHHC17 are indicated by arrows.

**Supplementary Table 1. Properties of each system, including the number of atoms and box sizes.**

| | Model | Total atoms | Protein atoms | Water | Ions | Box size, pre-npt (nm) | Box size, post-npt (nm) |
|---|---|---|---|---|---|---|---|
| wild-type | 0 | 89470 | 4301 | 28329 | CL:77; NA:81 | 12.26328, 9.32808, 7.84906 | 12.34600, 9.39100, 7.90200 |
| | 1 | 105617 | 4301 | 33702 | CL:91; NA:95 | 15.43770, 8.35268, 8.22443 | 15.52700, 8.40100, 8.27200 |
| | 10 | 134906 | 4302 | 43447 | CL:118; NA:121 | 12.24727, 11.82396, 9.36737 | 12.35400, 11.92700, 9.44900 |
| | 16 | 142490 | 4303 | 45971 | CL:124; NA:126 | 12.97844, 11.94914, 9.24683 | 13.02500, 11.99200, 9.28000 |
| mutant1 | 0 | 89126 | 4277 | 28223 | CL:76; NA:80 | 12.18691, 9.32384, 7.84862 | 12.27600, 9.39200, 7.90600 |
| | 1 | 105074 | 4277 | 33529 | CL:91; NA:95 | 15.36789, 8.34753, 8.22137 | 15.47000, 8.40300, 8.27600 |
| | 10 | 134505 | 4278 | 43322 | CL:117; NA:120 | 12.20899, 11.84468, 9.33716 | 12.29900, 11.93200, 9.40600 |
| | 16 | 141321 | 4279 | 45590 | CL:123; NA:125 | 12.90823, 11.93772, 9.23623 | 12.96800, 11.99300, 9.27900 |

**Supplementary Video 1. 250 ns molecular dynamics simulation of the ORF3–ZDHHC17 interaction**. The ZDHHC17 ARD (residues 55–288) is shown in blue and the ORF3 37–83 fragment in turquoise. The ORF3 PIFIQP motif and AAFIAA mutant (residues 58–63) are highlighted in red. ZDHHC17 amino acids within 0.4 nm of the PIFIQP motif or AAFIAA mutant are shown in orange. Hydrogen bonds are displayed as yellow dashed lines.
